# Cytosolic EZH2-IMPDH2 complex regulates melanoma progression and metastasis via GTP regulation

**DOI:** 10.1101/2021.11.02.467024

**Authors:** Gamze Kuser Abali, Fumihito Noguchi, Pacman Szeto, Youfang Zhang, Cheng Huang, Christopher K Barlow, Giovanna Pomilio, Christopher Chew, Samar Masoumi Moghaddam, Peinan Zhao, Miles Andrews, Isobel Leece, Jen G Cheung, Malaka Ameratunga, Nicholas C Wong, Ralf B Schittenhelm, Andrew Wei, Mark Shackleton

## Abstract

The enhancer of zeste homolog 2 (EZH2) oncoprotein is a histone methyltransferase that functions canonically as a catalytic subunit of the polycomb repressive complex 2 (PRC2) to tri-methylate histone H3 at Lys 27 (H3K27me3). Although targeting EZH2 methyltransferase is a promising therapeutic strategy against cancer, methyltransferase-independent oncogenic functions of EZH2 are described. Moreover, pharmacological EZH2 methyltransferase inhibition was only variably effective in pre-clinical and clinical studies, suggesting that targeting EZH2 methyltransferase alone may be insufficient. Here, we demonstrate a non-canonical mechanism of EZH2’s oncogenic activity characterized by interactions with inosine monophosphate dehydrogenase 2 (IMPDH2) and downstream promotion of guanosine-5’-triphosphate (*GTP*) production. EZH2-IMPDH2 interactions identified by Liquid Chromatography-Mass Spectrometry (LC-MS) of EZH2 immunoprecipitates from melanoma cells were verified to occur between the N-terminal EED-binding domain of cytosolic EZH2 and the CBS domain of IMPDH2 in a methyltransfersase-independent manner. EZH2 silencing reduced cellular GTP, ribosome biogenesis, RhoA-mediated actomyosin contractility and melanoma cell proliferation and invasion by impeding the activity of IMPDH2. Guanosine, which replenishes GTP, reversed these effects and thereby promoted invasive and clonogenic cell states even in EZH2 silenced cells. IMPDH2 silencing antagonized the proliferative and invasive effects of EZH2, also in a guanosine-reversible manner. In human melanomas, high cytosolic EZH2 and IMPDH2 expression were associated with nucleolar enlargement, a marker of ribosome biogenesis. EZH2-IMPDH2 complexes were also observed in a range of cancers in which Sappanone A (SA), which inhibits EZH2-IMPDH2 interactions, was anti-tumorigenic, although notably non-toxic in normal cells. These findings illuminate a previously unrecognized, non-canonical, methyltransferase-independent, and GTP-dependent mechanism by which EZH2 regulates tumorigenicity in melanoma and other cancers, opening new avenues for development of anti-EZH2 therapeutics.

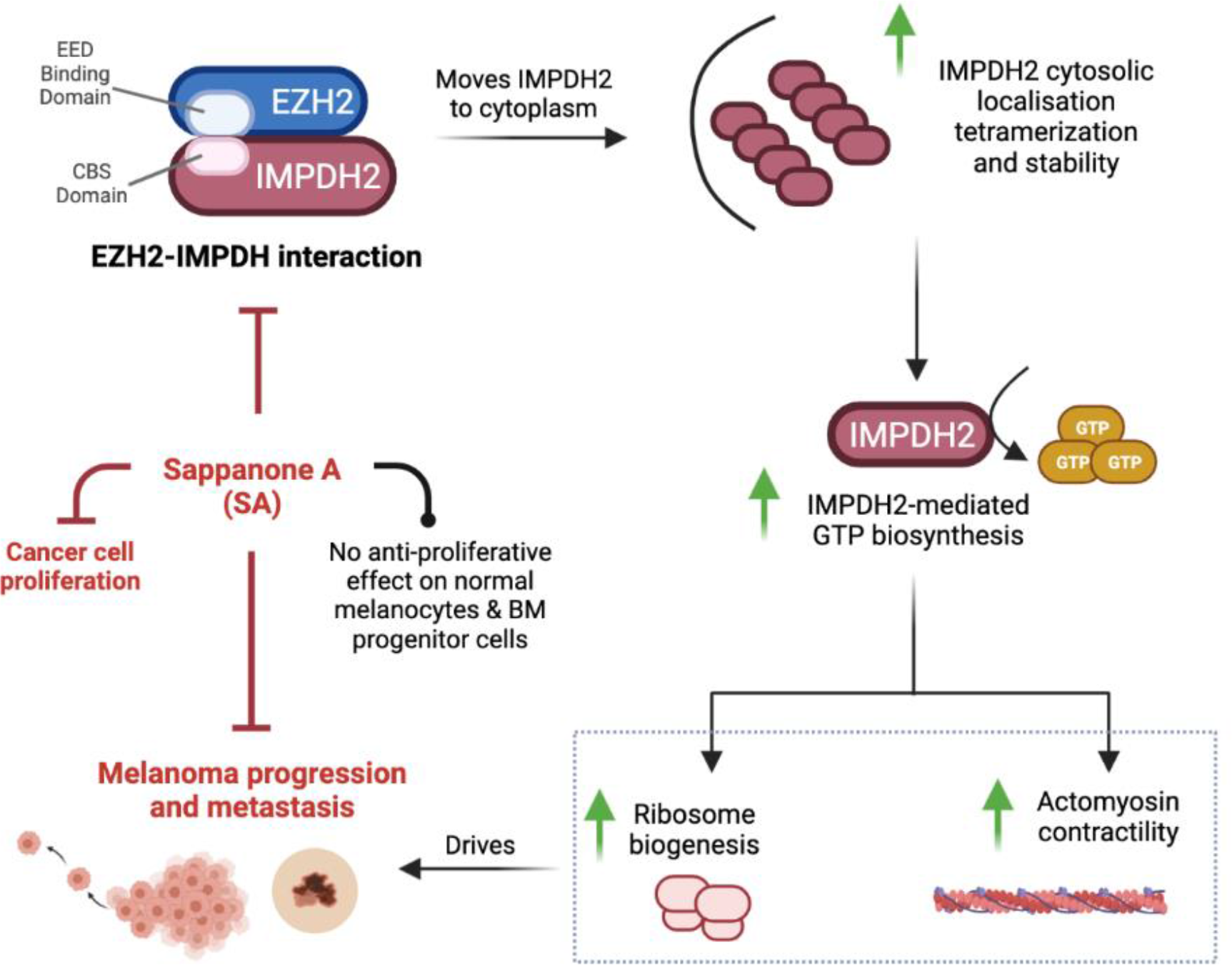
Graphical Abstract.

**Highlights:**

- EZH2 has non-canonical methyltransferase-independent and GTP-dependent tumorigenic and metastatic functions in melanoma.
- The N-terminal EED-binding domain of EZH2 interacts with the CBS domain of IMPDH2 in a polycomb repressive complex 2- (PRC2-) and methylation-independent manner.
- EZH2 accumulates with IMPDH2 in the cytoplasm and increases IMPDH2’s tetramerization-mediated activity independently of EZH2 methyltransferase.
- EZH2 upregulates GTP synthesis by IMPDH2 activation and thereby activates ribosome biogenesis via rRNA synthesis and actomyosin contractility via RhoA GTPase.
- Sappanone A (SA) inhibits IMPDH2-EZH2 interactions and is anti-proliferative across a range of cancers including melanoma, but not in normal cells.

## INTRODUCTION

Neoplastic cells, including melanoma, are highly dependent on *de novo* biosynthesis of purine nucleotides^1^. For example, the activity of Rho-GTPases in melanoma cells, and thereby formation of the actomyosin cytoskeleton which promotes cell migration and invasion, is regulated by intracellular GTP^2–4^. Consistent with this, cellular GTP levels, critical for purine nucleotide synthesis, are significantly higher in melanoma cells compared to their normal cell counterparts, melanocytes^2^.

Increased rRNA synthesis^5^ and nucleolar hypertrophy^6^ have long been recognized as features of malignant transformation. The requirement of GTP for Pol I transcription and nucleolar hypertrophy has been shown recently in glioblastoma^7^, and nucleolar hypertrophy has been associated with thicker and more mitotically active melanomas^8^. Selective inhibition of rRNA synthesis using the RNA polymerase I inhibitor CX-5461 decreased melanoma tumorigenicity *in vitro* and *in vivo*^9^.

Inosine monophosphate dehydrogenase 2 (IMPDH2), an oncoprotein in various cancers^10^, is a key rate-limiting enzyme in nucleotide synthesis. It maintains GTP levels needed for nucleic acid synthesis, protein production via ribosome biogenesis, and molecular signaling through guanine nucleotide-binding proteins (G-proteins) that regulate cell functions such as cytoskeletal rearrangement, membrane trafficking, and signal transduction^11^. IMPDH2 is regulated transcriptionally, post-translationally, and allosterically^12^, and tetramerization is essential for its activity^13, 14^. It contains two major domains: a catalytic domain for substrate interactions and the Bateman domain (CBS), which is not required for catalytic activity but exerts allosteric autoregulatory effects^13, 15, 16^. A naturally occurring compound, Sappanone A (SA), demonstrated inhibitory effects on neuroinflammation by directly targeting the conserved cysteine residue 140 (Cys140) in the CBS domain of IMPDH2. Interestingly, SA selectively targets and inactivates IMPDH2 but not the IMPDH1 isoform, potentially minimizing lymphotoxic effects of non-specific IMPDH family targeting^16^. IMPDH2 is overexpressed in melanoma cell lines compared to melanocytes^17, 18^, and depletion of GTP via IMPDH2 inhibition with MPA induced melanocytic differentiation in melanoma cells^19^.

Enhancer of zeste homolog 2 (EZH2), a component of Polycomb Repressor Complex 2 (PRC2), catalyzes tri-methylation of histone H3 at lysine 27 (H3K27me3) to regulate gene expression^20–22^. It has critical roles in the progression of numerous malignancies^23^, including melanoma^24–28^, where EZH2 activation represses tumor suppressor genes associated with cell differentiation, cell cycle inhibition, repression of metastasis, and antigen processing and presentation pathways^27–30^. EZH2 methyltransferase inhibitors have anti-cancer activity preclinically^31, 32^ and in patients ^31, 33^, albeit with notable toxicity.

Additional to EZH2’s methyltransferase activity, it also regulates gene transcription in a PRC2- and methylation-independent manner. This limits the therapeutic potential of specific EZH2 methyltransferase targeting^34^; compounds that degrade total EZH2 protein or that target methyltransferase-independent mechanisms of EZH2 might be required to avail context-dependent therapeutic potentials of EZH2 targeting.

We recently demonstrated that EZH2 is a negative regulator of melanocytic differentiation (pigmentation), whose suppression by knockdown or degraders decreased melanoma cell clonogenicity and invasion, and induced melanocytic differentiation^35^. In contrast, conventional EZH2 methyltransferase inhibitors displayed only minimal anti-melanoma efficacy *in vitro*^35^. These data further suggested methyltransferase-independent, non-catalytic functions of EZH2 in melanoma tumorigenesis and invasion, prompting us to look for novel EZH2 interactions.

Although EZH2 is mainly intranuclear, some studies have shown its cytosolic localization in fibroblasts, T lymphocytes, breast cancer, and prostate cancer cells^36–39^. Cytoplasmic functions of EZH2 are largely unknown, as most studies have focused on its nuclear functions. EZH2 is known to promote cancer progression by facilitating glucose, lipid, and amino acid metabolism^40, 41^, but other mechanisms of action are likely.

Here, we identify a previously unrecognized methyltransferase-independent role of EZH2 in melanoma tumorigenesis and invasion. We found that cytosolic EZH2 contributes to rRNA metabolism and Rho GTPase activity by regulating cytosolic IMPDH2 tetramerization-mediated activity and, in turn, promoting GTP production in melanoma cells. Sappanone A (SA) inhibited interactions between EZH2 and the IMPDH2 CBS domain and was anti-clonogenic in melanoma and a range of other cancer types, but not in normal cells. These findings suggest novel avenues for improved anti-EZH2 therapeutics.

## MATERIALS AND METHODS

### Mice

All animal experiments were performed in accordance with the Alfred Research Alliance Animal Ethics Committee protocols #E/1792/2018/M. All mice used in this study were supplied by and housed in AMREP Animal Services. Eight-week-old female NOD SCID. IL2R-/- Mice (NSG) mice were used for subcutaneous injection of pLV empty vector, shEZH2-3’UTR, shEZH2+EZH2-WT or shEZH2+ EZH2-H689A containing A375 melanoma cells (1x10^4^ cells mixed with 50 ul GFR-Matrigel, n=8 mice per group). Tumours were measured with callipers weekly, and all mice were sacrificed once the first tumour reached 20mm in diameter.

### Human Patient Samples

All human tissue related experiments, including human melanoma tissue FFPE samples, isolated primary human melanocytes and bone marrow samples were performed in accordance with the Alfred Human Research Ethics Committee protocols #155/18 and #29/05.

62 human melanoma tumor tissue sections ranging from grade I to IV were obtained from Melanoma Research Victoria (MRV) under the guidelines approved by the Victorian Government through the Victorian Cancer Agency Translational Research Program. MRV obtained the informed consent from all participants.

### Cell lines and primary cells

The HEK293, C32, SK-MEL28, IGR39, A375, B16-F10 and IGR37 cell lines were obtained from ATCC and cultured under conditions specified by the manufacturer. C006-M1 cell line was from QIMR Berghofer Medical Research Institute. LM-MEL28, LM-MEL33, LM-MEL43, LM-MEL45 were from Ludwig Institute for Cancer Research^42^. LM-MEL28: B4:F3 is the monoclonal line derived from LM-MEL28 cells in our lab previously. MCF7 and MDA-MB-231 cell lines were provided by Prof Jane Visvader (WEHI), OVCAR3, OVCAR8 cell lines were kindly provided by Prof David Bowtell (Peter MacCallum Cancer Centre), PC3, LNCaP, C4-2 cell lines by A Prof Renea A. Taylor (Monash Biomedicine Discovery Institute), OMM1 was kindly provided by Prof Bruce R. Ksander (Harvard Medical School) and 92.1 cell line by Prof Martine Jager (Leiden University Medical Centre). Mycoplasma tests were routinely performed in our laboratory and short tandem repeat (STR) profiling was conducted by the Australian Genome Research Facility (AGRF) to authenticate the human cell lines.

### Chemicals

The chemicals used for treating cells were GSK126 (Selleckchem, S7061), EPZ6438 (Selleckchem, S7128), Sappanone A (Cayman Chemicals, 23205), MPA (Selleckchem, S2487), Ribavirin (Selleckchem, S2504), DZNep (Sigma, S804983) and MS1943 (MedChemExpress, HY-133129); all are listed in Table S1.

### Plasmids, Cloning, Overexpression, and siRNA

pCMVHA hEZH2 (#24230) and pLV-EF1a-V5-LIC (#120247) plasmids were purchased from Addgene and MYC/FLAG-hIMPDH2 (#RC202977) plasmid from Origene. EZH2 (1-170), EZH2 (1-340), EZH2 (1-503), EZH2 (1-605), EZH2 (171-751) deletion mutants, full length EZH2 (1-751) and IMPDH2 (1-187) deletion mutant was cloned into pLV-EF1a-V5-LIC vector backbone’s SrfI/NotI RE using the cloning primers listed in Table S1. pCMVHA hEZH2 and V5-EZH2 vector was used to generate EZH2-H689A mutant vector using the mutagenesis primers listed in Table S1 with QuikChange II site-directed mutagenesis kit (Agilent) following the manufacturer’s instructions. Custom designed siRNA oligonucleotides listed in Table S1 were purchased from Bioneer Pacific. For transient transfection, 25x10^4^ cells were transfected with 2.5 µg of DNA using Lipofectamine 3000 transfection reagent (Invitrogen). For siRNA experiments, 25x10^4^ cells were transfected with 10 nM of the indicated oligonucleotides in Table S1 using the Lipofectamine RNAiMAX transfection reagent (Invitrogen). 72 hours after siRNA transfection, cells were used for functional assays or collected for western blot analysis.

Virus-containing supernatant was collected 48 hours after co-transfection of pCMV-VSV-G, psPAX2, pMD2.G and the EZH2 vectors into HEK293 cells, and then added to the target cells. Stable knockdown and rescue of EZH2 was achieved by lentiviral transduction of EZH2 with V5-EZH2-WT or V5-EZH2-H689A. After transduction, cells were selected for antibiotic resistance with 2 μg/mL puromycin (Sigma Aldrich, #P8833), followed by knockdown using stable short hairpin interfering RNA (MISSION shRNA, Sigma Aldrich) targeting the 3′UTR of human EZH2 (TRCN0000286227), as previously reported^38^.

### GST pull-down Assay

GST pull-down assay was performed as previously described^43^ with minor modifications. The plasmid GST-EZH2 (1-170), -EZH2 (1-340), -EZH2 (1-503), or – EZH2 (1-605) or GST only was expressed in BL-21 bacteria in the presence of 0.5mM IPTG for 2.5 h at 37°C. Bacterially expressed GST only (control) or each GST–EZH2 mutant peptide were solubilized in NETN buffer (1% NP-40, 20mM Tris-HCl, pH 8.0, 100mM NaCl, 1mM EDTA) and sonicated in 30 second bursts followed by 30 seconds rest for 15 cycles. Then they were purified by affinity chromatography on Glutathione Magnetic Agarose Beads (Pierce, Thermo Fischer)) and stored in PBS at 4°C until use. For GST–pull-down assays, purified GST control or GST–EZH2 mutant peptides were mixed with total lysates isolated from HEK293 cells, overexpressing V5-IMPDH2-CBS, grown in serum-fed condition and then incubated for 2 h at 4^°^C with constant rotation. The lysates from HEK293 cells were used as a source of IMPDH2-CBS domain. After extensive washing of unbound proteins, bound protein was eluted and analyzed by sodium dodecyl–PAGE (SDS–PAGE).

### CoImmunoprecipitation and HA/ FLAG pulldown assays

Pellets of 1x10^7^ cells were lysed with 1mL Co-IP Lysis Buffer (300mM NaCl, 50mM Tris HCL pH7.4, 0.5% NP40, 0.1% Sodium deoxycholate, 2% SDS) with PhosSTOP (Roche) and cOmplete (Roche) rolled at 4°C for one hour. DynaBeads™ Protein G (Thermofisher) were washed three times with Co-IP lysis buffer and chilled in preparation. Lysates were centrifuged at 15,000 RPM for 15 minutes at 4°C and the supernatant was collected, pre-cleared with 20µL of prewashed DynaBeads and incubated on a roller for 1 hour at 4°C. Lysates had pre-cleared beads removed and were split with 500µL for IgG control and 500µL for EZH2 sample, topped to 1 mL with Co-IP lysis Buffer. These were incubated overnight at 4°C with 1:1250 of Rabbit (DAIE) mAB IgG Isotype control or 1:300 of anti-EZH2 (D2C9) XP Rabbit antibody, respectively. After 16 hours of incubation, 35µL of pre-washed DynaBeads were added to IgG control or EZH2 sample and returned to the roller for 2-4 hours incubation at 4°C. Beads were washed with Co-IP buffer once, and then buffers of increasing salt concentrations (Buffer 1, 50mM Tris, pH8.0, 150mM NaCl; Buffer 2, 50mM Tris, pH8.0, 450mM NaCl; and buffer 3, 1M Tris, pH8.0). For Mass Spectrometry, proteins were eluted by resuspending in 150μL of 0.2M Glycine, pH2.5, for 5 minutes on ice and collecting supernatant, which was repeated twice. To the 450 μL of sample, 100 μL of 1M Tris-HCl (pH8.0) was added and the samples were kept at -80°C until LC-MS analysis. For CoIP-WB analysis beads were washed three times with Co-IP Lysis Buffer.

For HA pulldown assays the cells were lysed with 500 µL of IP Lysis Buffer containing cOmplete (Roche) protease inhibitor cocktail and incubated at 4°C for 35 min on a rotator followed by centrifugation at 15000 rpm for 15 min at 4°C. 50 µL of the lysates were kept for inputs. 25 µL of Pierce anti-HA magnetic beads (Thermo Fischer Scientific) were added onto the lysate and incubated at RT for 30 minutes on a rotator. Beads were washed with 300 µL of TBST three times and the beads were boiled in 2x SDS-Laemmli Sample Buffer for 10 minutes.

### Western blot

Total proteins were extracted from cell lines and tumor xenografts in ice-cold lysis buffer (10mM Tris HCL pH8.0 1mM EDTA 1% TritonX100 0.1% Sodium Deoxycholate 2% SDS 140mM NaCl, protease inhibitors and phosphatase inhibitors). Lysates were prepared after incubation on ice for 1h and centrifugation for 15 minutes cold at 15,000 rpm. Supernatants were boiled in 6x SDS-Laemmli Sample Buffer for 10 minutes. Proteins were run on 4–20% Mini-PROTEAN TGX Stain-Free Protein Gels (BioRad, 4568096) and then transferred to PVDF membrane by wet transfer system. Membranes were blocked with PBS containing 0.1% Tween-20 and 5% (w/v) skim milk, followed by incubation with the antibodies listed in Table S1. Signals were detected using Clarity ECL Western blotting Substrate (BioRad). Where applicable, signal intensities were quantified by ImageJ densitometry analysis software.

### Non-reducing SDS-PAGE

Samples were lysed in 2x Laemmli Sample Buffer without SDS and DTT and run with SDS free running buffer.

### DSS Crosslinking

1x10^6^ cells were precipitated and washed once with 1xPBS. The cell pellet was resuspended in 500 µL of 1xPBS. Cell suspensions were treated with either DMSO (control) or 1mM DSS (A39267, Thermo Fisher) and incubated for 30min at RT. Then, the cells were quenched with 50mM Tris-HCl pH:8.0 by incubating for 15min at RT. Finally, the cells were centrifuged, and the cell pellets were boiled in 50 µL of 2x SDS-Laemmli Sample Buffer for 10 min at 95^0^C.

### Cytoplasmic and nuclear fractionation

Cytoplasmic and nuclear extracts were isolated using a nuclear extraction kit according to the manufacturer’s protocol (Affymetrix; Santa Clara, CA) with modifications^43^. Co-IP was performed with anti-EZH2 or anti-IMPDH2 antibody at 4^°^C as described in the Co-IP method section. The immune complexes were collected with Protein G-Dynabeads (Thermo Fischer) and washed in lysis buffer. Bound proteins were analyzed by SDS–PAGE and WBs.

### RNA isolation and quantitative PCR

Total RNA from cells was extracted using Purelink RNA mini isolation kit according to the manufacturer’s instructions (Thermo Fischer Scientific) with the additional Purelink On-Column DNA purification (ThermoFisher Scientific) step. Complementary DNA (cDNA) was synthesized using total RNA (1 μg per reaction) with SuperScript Vilo cDNA synthesis kit (Thermo Fisher Scientific) as per manufacturer’s protocol. Quantitative PCR (qPCR) was carried out using Fast SYBR Green Master Mix (Invitrogen) and LightCycler 480 Instrument II (Roche). RNA expression changes were determined using a ΔΔCt method^44^. RPLP0 mRNA was used as an internal control in all qPCR reactions. Table S1 shows the qPCR primers used for IMPDH2, pre-rRNA, pre-tRNA, pre-GAPDH, p53, CDKN1A, CDKN2A, MDM2, PUMA and RPLP0 mRNA amplifications.

### Cell Proliferation, Clonogenicity and Sphere Formation Assay

To measure cell proliferation rates, we plated equal numbers of cells in 6-well plates. Cells were trypsinized and counted on the indicated days by haemocytometer after trypan blue staining.

For clonogenicity assay, cells were fixed with ice-cold absolute methanol for 20 min and air-dried for 15 minutes. Cells were stained with 0.5% Crystal Violet for 20 min at room temperature and then rinsed with tap water to remove excess dye. Five random fields of stained cells were imaged using bright field microscopy at 40 × magnification and average cell numbers per field were plotted as a function of time.

Sphere formation assays were performed as described^43^. Briefly, cells (500 cells per well) suspended in 100 mL ice-cold Matrigel in RPMI medium (1:1 ratio) were overlaid onto the pre-solidified 50% Matrigel in 24-well plates (100 ml per well). Cells were fed with 500 mL RPMI medium containing 10% FBS and grown for 14 days with a change of medium every 3 days. For the SA study, A375 and B16-F10 cells (500 cells/well) were grown on Matrigel and treated either with DMSO (vehicle) or SA in serum-fed conditions for the indicated days. Spheres were imaged and then manually quantified.

### Cell senescence β-Gal assay

Cells were fixed and stained with Senescence β-Galactosidase Staining Kit (CST #9860) following the manufacturer’s instructions and imaged on a Leica DMIL LED microscope under bright field settings.

### 3D Matrigel Invasion and Wound Healing Assays

Cells (1x10^5^ cells per well) were seeded in a 24-well Boyden chamber with an 8-mm filter coated with 20% growth factor reduced Matrigel. Cells were grown in RPMI medium containing 10% FBS for 16, 24 and 48 h at 37° C with 5% CO2. Cells on the inner side of the chamber were gently removed by scraping with a wet cotton swab. Invaded cells at the outer side of the chamber were fixed with 4% formaldehyde for 30 min at RT and rinsed twice with PBS. Cells were stained with 0.5% Crystal Violet for 20 min at RT and then rinsed with tap water to remove excess dye. Analysis was performed based on the average number of stained cells per field from f ive random fields at 20x magnification on a Leica DMIL LED microscope.

Wound healing assays were performed by seeding cells in complete media on a 24-well plate for 24–48 h until a confluent monolayer had formed. Linear scratches were made using a sterile 200 μl pipette tip. Monolayers were washed three times with PBS to remove detached cells, and then complete media was added. Images of the wound were taken immediately and 24h following wound formation on a Leica DMIL LED microscope under the phase contrast setting. Wound area was measured over time using ImageJ.

### RhoA activity assay

RhoA activities were measured in melanoma cells using RhoA G-LISA activation assay kit according to the manufacturer’s protocol. Briefly, cells were lysed in ice-cold lysis buffer and quickly cleared by centrifugation. Precision Red Advanced Protein Assay Reagent (Part # GL50) was used to quantify protein contents. Equal amounts of proteins were loaded onto ELISA plates. After several antibody incubation and washing steps the active RhoA bound protein levels were evaluated colorimetrically by OD490 nm absorbance measurement.

### IMPDH2 activity assay

IMPDH2 activity was measured by monitoring the reduction of NAD^+^ to NADH and the subsequent increase in absorbance at 340 nm in buffer: 100 mM Tris-HCl, 100 mM KCl, 2 mM DTT pH 7.4. 2 µg of recombinant IMPDH2 protein was preincubated with 3 mM IMP and 2 µg of recombinant EZH2 and then with 10 mM GTP for an additional 10 min before the reaction was initiated by the addition of 1 mM NAD^+^. NADH production was measured 1 h after incubation in a FLUOstar Omega plate reader by OD340 nm absorbance.

### Histochemical and immunostaining

H&E staining was done to evaluate nucleolar sizes^5^. For Fontana Masson staining, cells were sorted onto slides were fixed with 4% PFA for 20 minutes and washed twice with distilled water for 5 minutes. Then slides were incubated in Fontana silver nitrate working solution (2.5% Silver nitrate, 1% ammonium hydroxide) at 60^°^C for 2 hours. Slides were rinsed in water three times and incubated with 0.2% gold chloride solution (Sigma-Aldrich) for 2 minutes. Rinsed slides were incubated with 5% sodium thiosulfate for 2 minutes. After rinsing with water twice, slides were counterstained with 10 µg/mL DAPI solution for 5 minutes. Slides were rinsed and mounted with fluorescence mounting medium (Dako). For Schmorl’s staining, samples were dewaxed with 3 x 5-minute histolene washes, and rehydrated in washes of 100%, then 95% and then 75% ethanol for 5 minutes each. Slides were washed with distilled water and placed in Schmorl’s Stain for 10 minutes. Slides were washed with water for 1 minute, placed in Eosin in water for 15 seconds, and returned to constant washing with water for 3 minutes. The slides were finished with 4 x 2-minute washes of 100% ethanol and 3 x 2-minute washes of Histolene. Slides were mounted with DPX Mounting Medium (Thermo Fisher Scientific).

For melanoma patient sample IHC, slides were incubated at 60°C for 1h, dewaxed in histolene, and hydrated through graded alcohols and distilled water. Sections were subjected to antigen retrieval in Antigen Retrieval solution (Dako, pH6 for EZH2, IMPDH2 antibody) at 125° for 3 minutes heated by a pressure cooker. Primary antibody listed in Table S1 was diluted into blocking buffer and slides were incubated overnight at 4°C. After washing with TBST, the slides were incubated with secondary antibody using an ImmPRESS™ HRP Anti-Mouse IgG (Peroxidase) Polymer Detection Kit (Vector Laboratories) for 60 min at RT. Sections were washed with TBST and slides were developed by adding AEC+ High Sensitivity Substrate Chromogen Ready to use (Dako K346111-2).

For immunofluorescence, cells were fixed with 4% PFA diluted in PBS for 15 min at RT, rinsed three times with PBS, and blocked for 1h using blocking buffer (5% normal donkey serum containing 0.3% Triton X-100 in PBS). After blocking, slides were incubated with primary antibody (Table S1) diluted in antibody buffer (5% bovine serum albumin containing 0.3% Triton X100 in PBS) at 4 °C overnight. S lides were washed three times with PBS and incubated with fluorescent secondary antibodies indicated in Table S1. Slides were washed three times with PBS, stained with 10µg/ml DAPI and coverslipped using Fluorescence Mounting Medium (Dako). Slides were imaged using Leica DMIL LED inverted fluorescent microscope or Nikon A1r Plus si confocal microscope.

### Proximity ligation assays

Cells were seeded on round coverslips. After 24 h of seeding, cells were fixed with 4% PFA for 15 min at RT, rinsed three times with PBS, and blocked for 1 h using blocking buffer (5% normal goat serum containing 0.3% Triton X-100 in PBS). After blocking, slides were incubated with primary antibody diluted in antibody buffer (5% bovine serum albumin containing 0.3% Triton X-100 in PBS) at 4°C overnight. Slides were then washed three times with PBS and incubated with DuoLink PLA probes (Sigma, Cat #DUO92101). The protocol for PLA secondary antibody incubation, ligation, amplification, and washes were performed following the manufacturer’s protocol. Slides were imaged using a Nikon A1r Plus confocal microscope. Positive signals were normalized to single-primary antibody control (EZH2 or IMPDH2) and image analysis was performed using ImageJ.

### PDX Tumor Dissociation

Mice were euthanized with CO2 and tumors were resected. Tumors were manually dissociated in Hank’s Balanced Salt Solution (without Ca2+ and Mg2+, HBSS-/-), followed by enzymatic tumour dissociation using the gentleMACS tissue dissociator in Tissue Dissociating media (200 u/mL Collagenase IV, 5 mM CaCl2 in HBSS -/-). Tissue was washed with HBSS-/- and pelleted at 220g for 4 minutes at 4°C, and the supernatant was removed. After this, the pellet was resuspended with 100units/g of DNase and 5mL/g of warmed trypsinEDTA and incubated at 37°C for 2 minutes. Equal volumes of cold staining media were added, and the samples were pelleted at 220g for 4 minutes at 4°C. Supernatant was removed and the pellets were resuspended in cold staining media and filtered with a 40-micron cell strainer. To separate the tumoral cells from mouse stroma, cells were stained with an antibody cocktail of directly conjugated antibodies to mouseCD31 (endothelial cells), mouse CD45 (white blood cells), mouse TER119 (red blood cells) and human HLA-A/B antibodyin staining media on ice for 30 minutes. Labelled cells were resuspended in 2µg/ml DAPI in staining media with 10% FBS and 10uL/mL of DNase. Cells were subsequently analyzed and/or sorted on a FACSFusion (Becton Dickinson).

### CD34+ bone marrow progenitor cell isolation and culturing

Donor CD34+ HSPC samples were obtained from normal patients after informed consent in accordance with guidelines approved by The Alfred Health human research ethics committee. Cells from a leukapheresis sample were isolated using Ficoll-Paque PLUS (GE Healthcare) and density centrifugation, followed by NH4Cl lysis to remove red blood cells. A secondary isolation step was completed using CD34 MicroBead Kit (Miltenyi Biotec) performed according to the manufacturer’s protocol for positive selection of CD34+ cells from the mononuclear population. Isolated CD34+ cells were cultured in expansion medium (Stemspan SFEM (Stem Cell Technologies 09650), 50ng/mL rhFLt3L (R&D 308-FKN), 50ng/mL rhSCF (R&D 255-SC), 10ng/mL rhIL-3 (R&D 203-IL), 10ng/mL rhIL-6 (R&D 206-IL), 35nM UM171 and 500nM Stemreginin) with or without SA for 4 and 7 days.

### Human skin acquisition, single cell suspension, isolation of melanocytes via FACS and melanocyte culture

Epidermal melanocytes were isolated from normal adult human breast skin. The skin samples were provided from Caucasoid donors (age 18 – 72) via The Victorian Cancer Biobank. Fat was removed from the skin and washed in PBS with Gentamycin (10 µg/mL) and 80% EtOH. Then the skin was cut into small pieces (∼5 mm^2^) and incubated in Dispase (15 U/mL, Gibco/Thermo Fisher Scientific) with Gentamycin (10 µg/mL) at 4°C overnight. Epidermis was peeled from dermis by forceps and smashed by scissors and incubated in Trypsin/EDTA (0.25%) at 37°C for 10 min to make a single cell suspension of the epidermal cells. After pipetting and addition of fetal bovine serum (FBS) to stop activity of trypsin (final concentration of FBS is 10%), the epidermal single cell suspension was passed through cell-strainers (70 µm then 40 µm). After centrifugation (220g for 5 min) the collected epidermal cells were suspended in the staining medium and viability was validated microscopically with trypan blue.

The collected epidermal cells from skin were incubated with primary antibodies including FITC anti-human CD326 (EpCAM) (1:100, mouse), FITC anti-human CD31 (1:100), FITC anti-human CD45 (1:100), FITC anti-human CD235a (1:100) and PE anti-human CD117 (c-kit) (1:100) in the staining media for 30 min at 4°C. After a wash and centrifuge the cells were suspended in DAPI (2.5 µg/mL) and subjected to FACS analysis (BD FACSAria™ Fusion flow cytometer, BD). Debris (by morphology plot: FSC-A/SSC-A), doublets (by doublet plot: FSC-H/FSC-W and SSC-H/SSC-W) and dead cells (DAPI+) were excluded. The Ckit+CD326-CD31-CD45-CD235a-fraction was sorted into 1.5 mL microcentrifuge tubes filled with Medium 254 (1 mL).

The sorted primary human melanocytes were plated on HaCaT-derived ECM-coated culture dish and cultured in Medium 254 supplemented with Human Melanocyte Growth Supplement-2 (HMGS-2, including basic FGF, insulin, transferrin, bovine pituitary extract, endothelin-1, FBS, heparin and hydrocortisone, concentrations are proprietary, PMA-free) at 37°C with 10% O2 and 5% CO2.

### RNA-Seq data analysis

FASTQ files were processed using Laxy (https://zenodo.org/record/3767372) which encompasses the RNAsik pipeline (https://joss.theoj.org/papers/10.21105/joss.00583). Briefly, GRCh38 reference genome was used for STAR alignment^45^ and gene expression counts were performed using featureCounts^46^. Gene counts were analysed using Degust (https://zenodo.org/record/3501067) for differential expression analysis. Data processing was performed on NeCTAR Cloud Servers, or MASSIVE High Performance Computing (HPC) cluster.

Differential gene expression analysis was performed using edgeR (v.3.32.1). Quasi-likelihood F-test was performed with glmQLFit and glmQLFTest functions. Gene ontology (GO) enrichment test was performed using PANTHER (v16.0) fisher’s exact test corrected by false discovery rate (FDR).

### TCGA survival analysis

The clinical data and mRNA expression profiles for skin cutaneous melanoma samples in TCGA PanCancer Atlas database were retrieved from MSKCC Cancer Genomics Data Server (CGDS) (http://www.cbioportal.org)47. The “high expression” and “low expression” groups for each gene were defined as above or below the median expression level for the cohort respectively. The overall survival (OS) curves were calculated with the Kaplan-Meier method and the statistical significance were tested with the log-rank test. The calculations were performed using the R package ‘survival’ 3.1-11 and the survival curves were plotted using the R package ‘survminer’ 0.4.4.

### Sample preparation for GTP analysis

1 x10^7^ A375 cells were washed once with 0.9% NaCl and cell pellets were snap frozen prior to LC-MS analysis. 200 µL of extraction solvent (2:6:1 CHCl3: MeOH: H2O) at 4°C was added to the washed cell pellets after which the samples were briefly vortexed, sonicated in an ice-water bath (10 minutes). Samples were then frozen in liquid nitrogen and thawed three times before mixing on a vibrating mixer at 4°C for 10 minutes after which they were subjected to centrifugation (20,000 x g, 4°C, 10 min) and the supernatant transferred to samples vials for prompt (same day) LC-MS analysis.

### LC−MS analysis for metabolomics

Samples were analyzed by hydrophilic interaction liquid chromatography coupled to triple quadrupole mass spectrometry (LC−MS). In brief, the chromatography utilized a ZIC-p(HILIC) column (Merck SeQuant ZIC-pHILIC 5um 150 x 4.6 mm, polymeric) and guard (Merck SeQuant ZIC-pHILIC Guard, 20 x 2.1 mm, PEEK coated guard) with a gradient elution of 20 mM ammonium carbonate (A) and acetonitrile (B) (linear gradient time-%B as follows: 0 min-80%, 15 min-50%, 18 min-5%, 21 min-5%, 24 min-80%, 32 min-80%) on a 1290 Infinity II (Agilent). The flow rate was maintained at 300 μL/min and the column temperature 25°C. Samples were kept at 10°C in the autosampler and 5 μL injected for analysis. The mass spectrometry was performed in multiple reaction monitoring (MRM) mode on an Agilent 6495 Triple Quadrupole. Full details are provided in supplementary material. Peak integration was carried out using MassHunter Qualitative Navigator B.08.00 (Agilent).

### LC-MS analysis for proteomics

Immunoprecipitated proteins were reduced with 10 mM TCEP (Thermo Fisher), alkylated with 40 mM iodoacetamide (Sigma Aldrich) and digested with sequencing grade trypsin (Promega). Samples were acidified with 1% formic acid (FA) and purified using OMIX C18 Mini-Bed tips (Agilent) prior to LC-MS/MS analysis.

Using a Dionex UltiMate 3000 RSLCnano system equipped with a Dionex UltiMate 3000 RS autosampler, an Acclaim PepMap RSLC analytical column (75 µm x 50 cm, nanoViper, C18, 2 µm, 100Å; Thermo Scientific) and an Acclaim PepMap 100 trap column (100 µm x 2 cm, nanoViper, C18, 5 µm, 100Å; Thermo Scientific), the tryptic peptides were separated by increasing concentrations of 80% acetonitrile (ACN) / 0.1% formic acid at a flow of 250 nl/min for 128 min and analyzed with a QExactive HF mass spectrometer (ThermoFisher Scientific). The instrument was operated in the data dependent acquisition mode to automatically switch between full scan MS and MS/MS acquisition. Each survey full scan (m/z 375–1575) was acquired in the Orbitrap with 60,000 resolution (at m/z 200) after accumulation of ions to a 3 x 10^6^ target value with maximum injection time of 54 ms. Dynamic exclusion was set to 15 seconds. The 12 most intense multiply charged ions (z ≥ 2) were sequentially isolated and fragmented in the collision cell by higher-energy collisional dissociation (HCD) with a fixed injection time of 54 ms, 30,000 resolution and automatic gain control (AGC) target of 2 x 10^5^.

Raw data files were analyzed with the MaxQuant software suite v1.6.5.0^48^ and its implemented Andromeda search engine^49^ to obtain protein identifications and their respective label-free quantification (LFQ) values using standard parameters. The proteomics data were further analyzed using either Perseus^50^ or LFQ-Analyst^51^.

### Statistical Analysis

Analysis was performed using GraphPad Prism version 8. All analyses were performed using log-rank test, unpaired two-tailed t-tests, one-way or two-way ANOVA plus Tukey’s multiple comparison tests as appropriate to the data type. The statistical parameters are reported in figure legends or text of the results. P values less than 0.05 were considered significant.

## RESULTS

### Methyltransferase-independent activity of EZH2 in melanoma

We recently reported that decreasing EZH2 abundance rather than EZH2 methyltransferase activity may be a key to realizing the therapeutic potential of EZH2 targeting in melanoma^35^. Further to investigate methyltransferase-independent functions of EZH2 in melanoma, we compared melanoma cells subjected to EZH2 targeting by siEZH2 knockdown, or treatment with the EZH2 degrader DZNep or with EZH2-methyltransferase inhibitors GSK126 and EPZ-6438.

Although GSK126 and EPZ-6438 inhibited EZH2 methyltransferase activity as measured by H3K27me3 levels (Fig. 1A, S1A and S1B), they had no effect on the growth, clonogenicity, migration, invasion, or pigmentation of BRAFV600E mutant A375 and IGR37 melanoma cells, and only partial effects on NRASQ61K mutant C006-M1 cells (Fig. 1A-1F and S1A-S1H). In contrast, EZH2 knockdown or DZNep treatment displayed marked anti-melanoma effects and promoted melanocytic differentiation in all lines tested (Figure 1A-D and S1A-1H). These findings provide further evidence of methyltransferase independent functions of EZH2 in melanoma.

**Figure 1.**
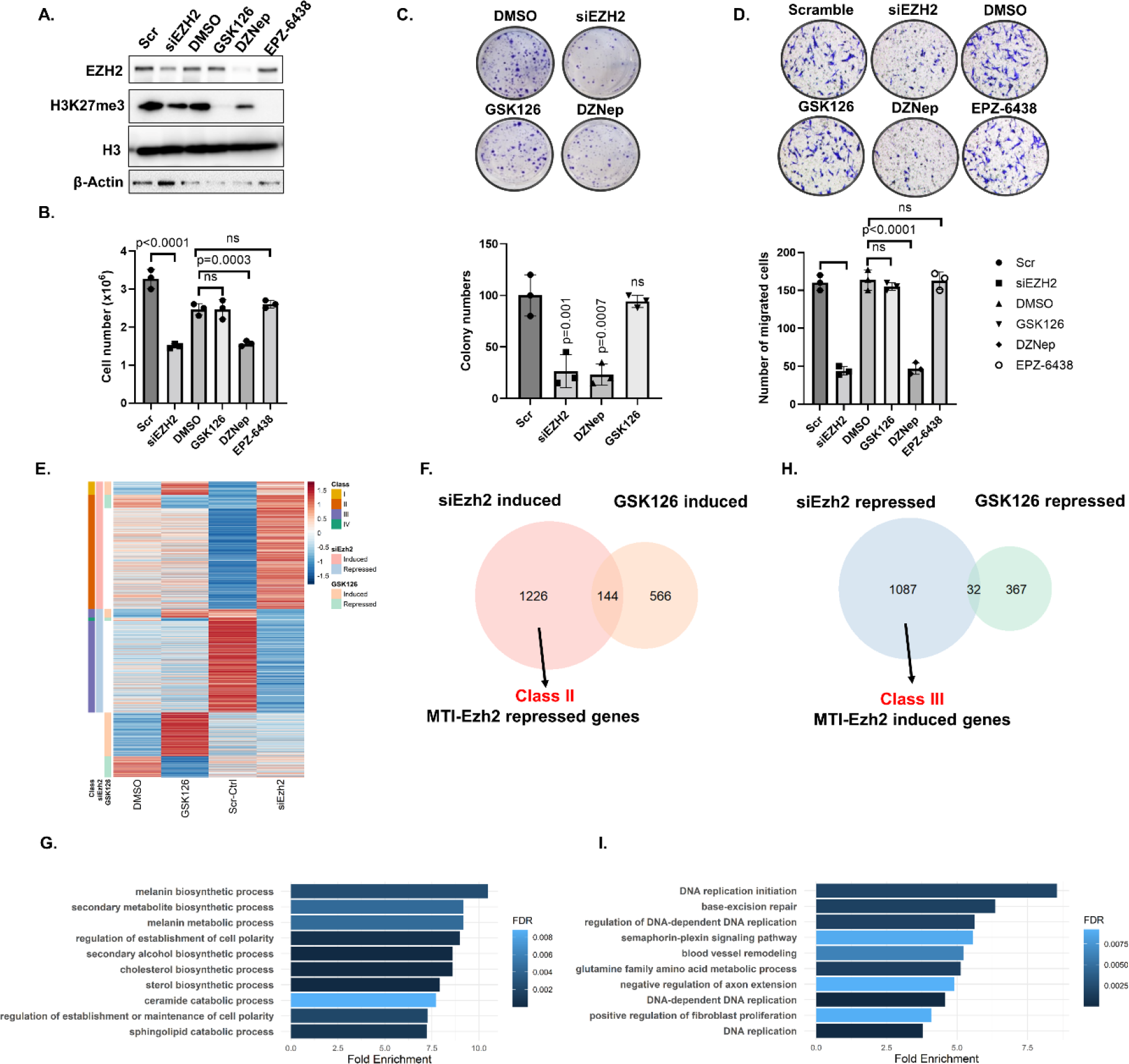
Methylation-independent functions of EZH2 are predominant in melanoma. A375 cells were treated with siEZH2, 2 μM DZNep, 2 μM GSK126, 2 μM EPZ6438 and scramble or DMSO (control) for 3 days prior to: (A) Western blot analysis of EZH2, H3K27me3, H3 and β-Actin protein level, (B) cell growth analysis done by Trypan Blue haemocytometer counting, (C) clonogenicity after low-density seeding (crystal violet stain). Clonogenicity was assessed in pre-treated (3 days) cells seeded at 2000 cells in 6-well plate followed by crystal violet staining (0.5% in methanol) after incubation for 10 days in drug-free media. Representative images after crystal violet-stained wells were shown above bars (D) Boyden chamber migration was assessed in pre-treated (3 days) cells seeded at 50,000 cells in 24-well plate after incubation for 24h. Representative images after crystal violet-stained wells were shown above bars. (E) B16-F10 cells were treated with either GSK126 versus vehicle control (DMSO) or siEzh2 versus siCtrl and then profiled in triplicate RNAseq experiments. Genes that were significantly up- or downregulated by siEzh2 compared with the control were clustered across all samples and are shown as heatmaps. Each row represents one gene and each column triplicate sample. The siEzh2-induced genes that were also induced by GSK126 were termed class I genes and those unchanged by GSK126 class II genes. Genes that were activated by Ezh2 were defined as class III genes. (F) Venn diagram showing overlap among si-Ezh2 induced and GSK126-induced genes compiled from RNAseq experiment in G. (G) GO biological process analysis of 1226 class II genes. (H) Venn diagram showing overlap among si-Ezh2 repressed and GSK126-repressed genes compiled from RNAseq experiment in G. (I) GO biological process analysis of 1087 class III genes. Data for B-D are from three independent experiments and are presented as mean ±SD, analyzed by one-way ANOVA plus Tukey’s multiple comparison tests. ns: non-significant.

To examine methyltransferase-dependent and -independent transcriptional programs of EZH2, we performed global gene expression analysis in B16-F10 murine melanoma cells treated with siEzh2 knockdown vs siRNA control, and also with GSK126 vs DMSO control. 1370 genes were significantly increased by siEzh2 depletion (Figure 1E, 1F), of which 1226 (89.5%) were not upregulated by GSK126 treatment (Figure 1F). By gene ontology (GO) analysis, these 1226 genes were enriched in melanin and cholesterol biosynthesis pathways (Figure 1G). Of the 1119 genes that were downregulated by siEzh2 (Figure 1E-1H), 1087 (97.1 %) were not changed upon GSK126 treatment (Figure 1H) and strongly enriched in DNA replication and DNA repair pathways (Figure 1I). These data are consistent with regulation by EZH2 of methyltransferase-dependent as well as -independent transcriptional programs in melanoma.

Because siRNA might have off-target effects, we also tested a catalytically dead mutant of EZH2, H689A, which lacks methyltransferase activity. A375 cells were treated with control (sh-control) or 3’ UTR EZH2 region-targeting shRNA (Figure 2A and S2A) to deplete endogenous EZH2, which was then rescued with either wild-type (V5-EZH2-WT) or H689A-mutant EZH2 (V5-EZH2-H689A). *In vitro* and *in vivo*, shEZH2 3′UTR knockdown reduced A375 and IGR37 clonogenicity, invasion, wound healing and tumour formation, and increased pigmentation, in a manner that was reversed similarly by ectopic expression of V5-EZH2-WT and V5-EZH2-H689A (Figure 2B-F and S2B-D).

**Figure 2.**
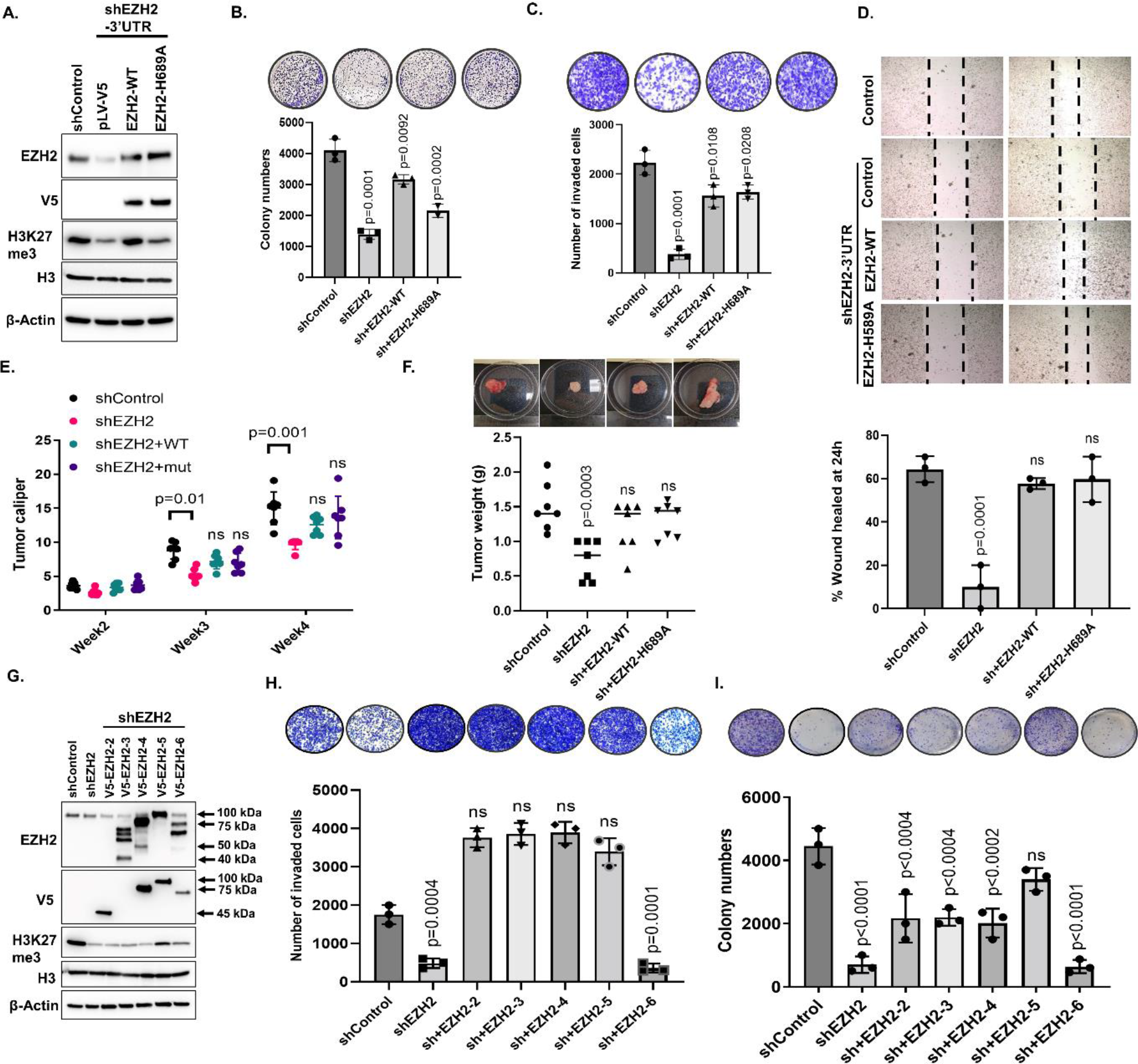
EZH2 has methyltransferase independent function in melanoma tumorigenicity and invasion. (A) Western blot analysis of A375 cells showing EZH2 knockdown after lentiviral transduction with control shRNA (shControl) or 3′ UTR EZH2-targeting shRNA (shEZH2) and rescue with V5-tagged WT-EZH2 or methyltransferase deficient H689A-EZH2. (B) Clonogenicity assay of cells described in A. Representative images after crystal violet-stained wells were shown above bars. (C) Matrigel-coated Boyden chamber invasion assay of cells described in A. Representative images after crystal violet staining were shown above bars. (D) Wound healing assay of cells described in A. Representative images of the wound after 24 h shown above bars. (E) Tumor caliper of A375 xenografts as described in A. (F) Tumor weights of A375 xenografts at the end point. Representative tumors per group were shown above bars. (G) Western blot analysis, (H) invasion, (I) clonogenicity of A375 cells with EZH2 knockdown followed by rescue with V5-tagged EZH2 deletion mutant vectors. Data for B-D, H, I are from at least three independent experiments and are presented as mean ±SD, analyzed by one-way ANOVA plus Tukey’s multiple comparison tests. Data for E, F are from 7 mice per group and are presented as mean ±SD, analyzed by two-way ANOVA plus Tukey’s multiple comparison tests. ns: non-significant.

To complement these data, we generated deletion mutants to interrogate EZH2 domains that might rescue shEZH2 knockdown phenotypes (Figure 2G). Partial rescue of A375 cell clonogenicity and full rescue of cell invasion were observed in all shEZH2 3′UTR knockdown cells co-transfected with EZH2 deletion constructs that lacked the SET domain, which encodes methyltransferase. In contrast, rescue was not consistently observed with N-terminal EED domain deletion mutants (Δ1-169) [clone 6] with intact SET domains (Figure S2E, Figure 2G-I), confirming that the N-terminal EED domain of EZH2, rather than the methyltransferase-containing SET domain, is critical for the clonogenicity and invasion in melanoma.

### Interactions between the EZH2 and IMPDH2

To characterize methyltransferase-independent actions of EZH2, we examined its interacting partners in melanoma by immunoprecipitating EZH2 from protein lysates and subjecting the precipitate to liquid chromatography mass spectrometry (LC-MS). Expected PRC2 complex proteins were identified as EZH2 binding partners in four *BRAF^V^*^600^*^E^* mutated cell lines, one *NRAS^Q61K^* mutated line, and in one *BRAF^V^*^600^*^E^* PDX melanoma (Figure 3A and Table S2). Additionally, ubiquitin degradation pathway proteins UBR4 and NPLOC4, Kinesin 1 complex proteins including KIF5B, KLC1, KLC2 and KLC4, and Inosine-5’-monophosphate dehydrogenase 2 (IMPDH2), were all consistently co-immunoprecipitated with EZH2.

**Figure 3.**
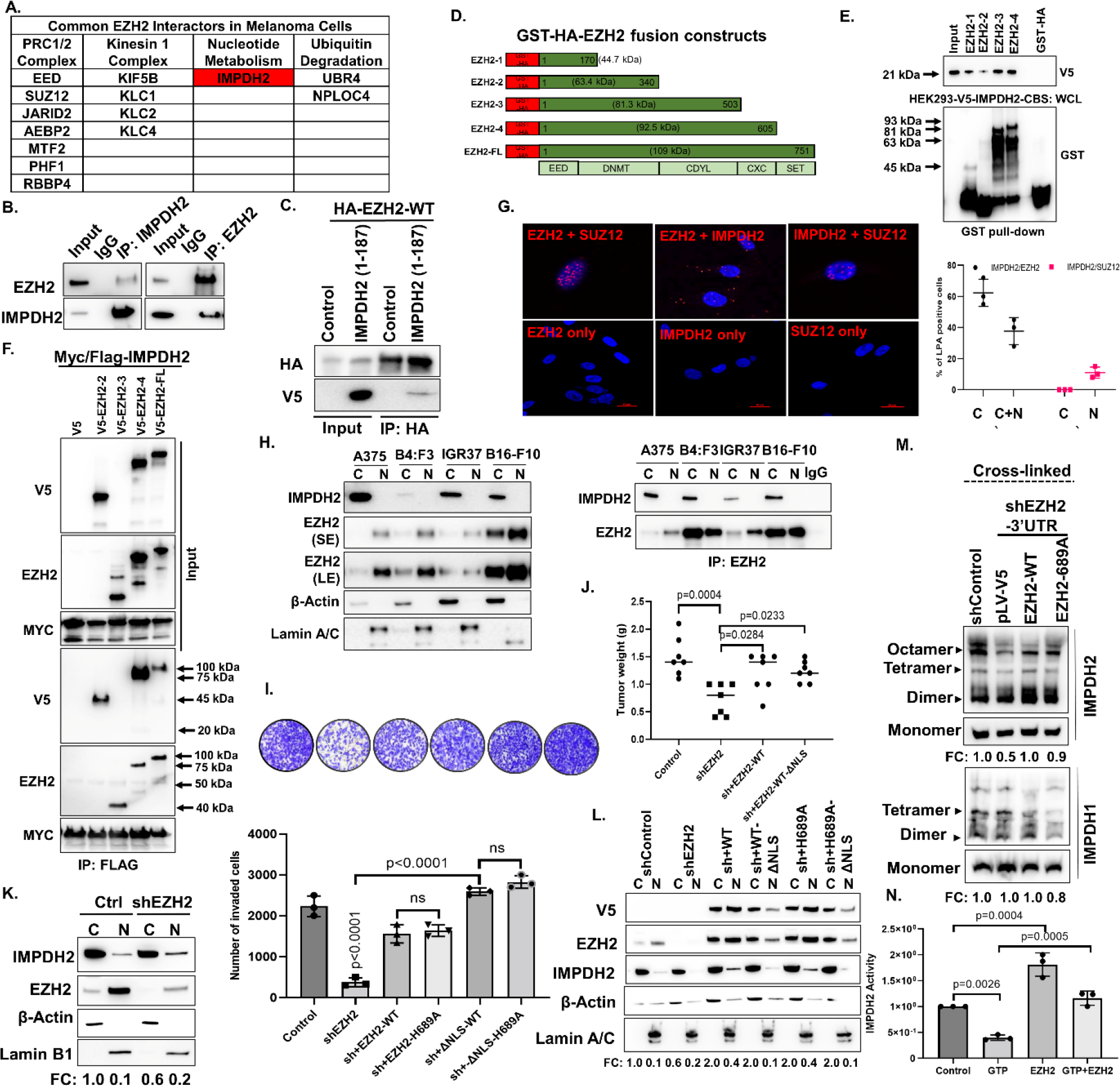
Cytosolic EZH2 interacts with IMPDH2 through the IMPDH2-CBS domain and moves IMPDH2 to cytoplasm/ increases its tetramerization-mediated activity. A) List of overlapping proteins co-immunoprecipitated (Co-IP) with EZH2 from C006-M1, LM-MEL-28:B4:F3, IGR37, A375 and LM-MEL-45 melanoma cells (all data derived from n=3 biological replicates). (B) The interaction between endogenous EZH2 and IMPDH2 was determined in A375 cells by immunoprecipitation (IP) with anti-IMPDH2 and anti-EZH2 antibody followed by western blotting with anti-EZH2 and anti-IMPDH2 antibody. (C) HA-tagged EZH2-WT and V5-tagged IMPDH2 (1-187) were co-expressed in A375 cells. The interaction between overexpressed EZH2 and IMPDH2 (1-187) was determined by immunoprecipitation with anti-HA antibody followed by western blotting with anti-V5 antibody. (D) GST-EZH2 deletion mutant constructs. (E) The binding of V5-IMPDH2-CBS protein to GST–EZH2 peptides was probed with WB using the V5 specific antibody. Total cell lysate from HEK293 overexpressing V5-IMPDH2-CBS was used as a source of IMPDH2-CBS in GST–pull-down experiment. (F) The binding of Myc/Flag tagged full length IMPDH2 protein to V5 tagged EZH2 deletion mutant peptides (shown in Fig. 2G) was shown by co-IP with anti-Flag antibody followed by probing with anti-V5 antibody. (G) Ligation proximity images depicting co-localization with EZH2 and IMPDH2 by red fluorescent dots A375 cells. Scale bar=10 μm. Number of interaction loci depicted as red dots were counted for cytoplasm and nucleus of total of 100 cell. (H) Cytosolic/Nuclear fractionation was done for A375, B4:F3, IGR37, B16-F10 cells followed by IP with anti-EZH2 antibody followed by western blotting with anti-EZH2 and anti-IMPDH2 antibody (right). Lamin A/C is nuclear, and β-Actin is cytosolic marker. Inputs were shown on the left. (I) Matrigel-coated Boyden chamber invasion assay of A375 cells showing EZH2 knockdown after lentiviral transduction with control shRNA (shControl) or 3′ UTR EZH2-targeting shRNA (shEZH2) and rescue with V5-WT-EZH2 or V5-EZH2-H689A, V5-EZH2-ΔNLS-WT and V5-EZH2-ΔNLS-H689A. Representative images after crystal violet staining were shown above bars. (J) Tumor weights of indicated A375 xenografts (n=7) at the end point. (K) Cytosolic/Nuclear fractionation was done from A375 cells with control shRNA (shControl) or 3′ UTR EZH2-targeting shRNA followed by IP with anti-EZH2 antibody followed by western blotting with anti-EZH2 and anti-IMPDH2 antibody. Lamin A/C is nuclear, and β-Actin is cytosolic marker. (L) Cytosolic/Nuclear fractionation was done from cells described in (I) followed by western blotting with anti-V5, anti-EZH2, anti-IMPDH2 antibody and β-Actin antibody. (M) The clusters of IMPDH2 and IMPDH1 tetramer were detected from cross-linked whole-cell extracts isolated from cells described in Figure 2A. (N) Relative IMPDH2 activity measured by NADH absorbance at OD340nm. 2 µg of recombinant IMPDH2 was preincubated with and 3 mM IMP and 2 µg of recombinant EZH2 and then 10mM GTP was added for 10 min prior to reaction initiation by with 1 mM NAD^+^. Data for I, J and N is from three independent experiments and are presented as mean ±SD, analyzed by one-way ANOVA plus Tukey’s multiple comparison tests. ns: non-significant.

We focused on IMPDH2-EZH2 interactions because of IMPDH2’s known role in GTP metabolism^52^. Further, we found that IMPDH2 protein level is upregulated in human melanoma cells compared to normal human melanocytes (Figure S3A) and that high IMPDH2 expression is correlated with poor melanoma survival (p= 0.01, cbioportal) (Figure S3B). To investigate EZH2-IMPDH2 interactions, we verified them endogenously in lysates from A375 cells and PDX tumors by reciprocal Co-IP (Figure 3B and S3C), finding. Interactions were reproducibly seen even after GSK126 or EPZ6438 treatment, suggesting they occur independently of EZH2 methyltransferase activity (Figure 3SD). Furthermore, we were unable to detect IMPDH2 methylation in EZH2 Co-IPs by mass spectrometry (Table S3).

To define interacting domains, we exogenously co-expressed tagged EZH2 (HA-EZH2) with both full length IMPDH2 (MYC/FLAG-IMPDH2) in HEK293 cells (Figure S3E) and with the CBS domain of IMPDH2 [V5-IMPDH2-(1-187)] in A375 cells (Figure 3C). GST pull down assays showed that the CBS domain of IMPDH2 interacts with the N-terminal EED binding domain (1-170) of EZH2 (Figure 3D, 3E). EZH2 EED domain interactions with full length IMPDH2 were verified using exogenously expressed V5-EZH2 deletion mutants and MYC/FLAG-IMPDH2 (Figure 3F).

Further to characterize EZH2-IMPDH2 interactions, we also performed proximity ligation assays (PLAs). 60% of A375 cells showed cytosolic EZH2-IMPDH2 interactions (<40nm apart, average 15 foci per cell), and 40% showed both cytosolic and nuclear interactions (average 4 foci per cell were nuclear, Figure 3G). These results were supported by western blotting of separated cytosolic and nuclear protein fractions (Figure 3H) and multiplex immunofluorescence labelling of melanoma cell lines (Figure S3F). In contrast, we did not detect cytosolic IMPDH2 interactions with the PRC2 component SUZ12 by PLA, although 10% of cells showed PLA positive nuclear foci (average 4 per cell, Figure 3G).

These data indicate that although EZH2 is mostly localized in nuclei, its N-terminal EED domain interacts directly with the IMPDH2-CBS domain predominantly in the cytosol and independently of PRC2 complex formation or EZH2 methyltransferase activity in melanoma cells.

### Cytosolic EZH2 drives melanoma progression

To test a potential role for cytoplasmic EZH2 in melanoma progression, we developed an EZH2 mutant lacking a nuclear localization domain (EZH2-ΔNLS). We first generated A375 cells with stable 3′UTR EZH2 knockdown (to minimize endogenous EZH2) and then rescued EZH2 with full length (V5-EZH2-WT) or V5-EZH2-WT-ΔNLS lentiviral constructs. V5-EZH2-WT-ΔNLS expression was mostly cytoplasmic and depleted nuclear EZH2 methyltransferase activity on histone H3K27 (Figure S3G).

A375 shEZH2 melanoma cells and xenograft tumors displayed reduced invasion and tumorigenicity that were restored similarly by WT-EZH2 and V5-EZH2-WT-ΔNLS (Figure 3I, 3J and S3H). Moreover, overexpression of cytosolic EZH2 lacking methyltransferase activity (V5-EZH2-H689A-ΔNLS) also restored the invasive phenotype of shEZH2 A375 cells to levels comparable to those achieved with V5-EZH2-WT-ΔNLS. Although p38-dependent phosphorylation of EZH2 at its T367 residue was shown to induce cytosolic localization of EZH2 in breast cancer^38^, analysis of post-translational modifications of EZH2 in our EZH2-IP LC-MS data did not show significantly different phosphorylation isoforms between cytosolic and nuclear EZH2 (Figure S3I, Table S4). Thus, cytoplasmic EZH2 expression is sufficient to promote melanoma cell invasion and tumorigenicity irrespective of EZH2 nuclear or methyltransferase function.

### Increased cytosolic localization and activation of IMPDH2 by EZH2

We next investigated effects of EZH2-IMPDH2 interactions on IMPDH2 localization and tetramerization/activity. Stable EZH2 knockdown slightly decreased total IMPDH2 protein, but not mRNA expression, that was later rescued by overexpression of EZH2 (1-340) [clone 2] (Figure S3J). Fractionation and immunofluorescence experiments showed that stable or transient EZH2 knockdown increased the nuclear localization of endogenous IMPDH2 and exogenously expressed IMPDH2-CBS domain (Figure 3K, S3K, S3L). Conversely, overexpression of cytosolic wild-type EZH2 (V5-EZH2-WT-ΔNLS) or of cytosolic EZH2 (1-340) [clone 2] increased cytosolic IMPDH2 compared to overexpression of wild type EZH2 (V5-EZH2-WT) in endogenous EZH2-silenced A375 cells (Figure 3L, S3M), independently of EZH2 methyltransferase activity (Figure 3L and S3G).

As tetramerization is an essential step in IMPDH2 activation ^13, 16, 53^, we investigated effects of EZH2 on IMPDH2 tetramerization and activation in A375 cells. Cross-linked whole-cell extracts showed that IMPDH2 rather than IMPDH1 tetramers were decreased by stable/transient EZH2 knockdown that was rescued by overexpression with wild-type or methyltransferase-deficient EZH2 (full length and EZH2 (1-340) [clone 2], Figures 3M, S3N and S3O). Additionally, co-incubation of IMPDH2 with EZH2 increased basal IMPDH2 activity and reversed GTP-mediated allosteric inhibition of IMPDH2 (Figure 3N).

### GTP-dependent regulation by IMPDH2 of ribosome biogenesis and actomyosin contractility

We next assessed pharmacological and genetic inhibition of IMPDH2 in melanoma. Treatment with mycophenolic acid (MPA) or ribavirin, pan-IMPDH inhibitors^12, 54, 55^, decreased cell proliferation, clonogenicity and invasion that was rescued by guanosine addition regardless of B-Raf or N-Ras mutational status (Figure 4A-E and 4A-S4E). MPA and ribavirin also induced pigmentation and senescence (Figure S4F, S4G). siRNA silencing of IMPDH2 also retarded melanoma cell proliferation and invasion that was restored by guanosine addition (Figure 4F, 4G, S4H and S4I). These results implicate a GTP-dependent role for IMPDH2 in melanoma progression.

**Figure 4.**
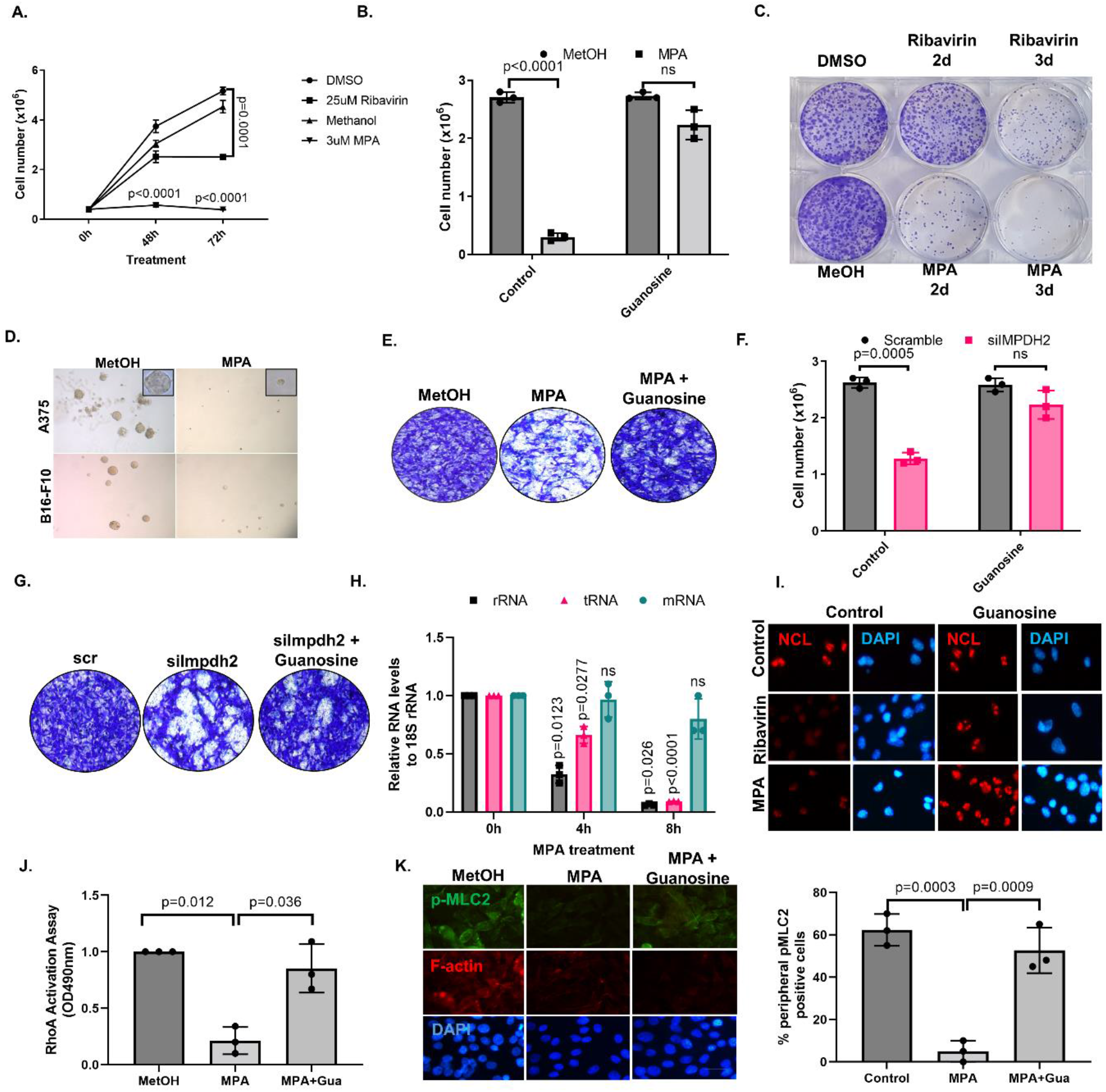
IMPDH2 induces clonogenecity/ invasion in melanoma cells by regulating ribosome biogenesis and actomyosin contractility via cellular GTP level regulation. (A) Time-dependent growth curves of A375 cells upon 25 µM Ribavirin or DMSO (control); 3 µM MPA, or methanol (Control). (B) A375 cell growth analysis done by Trypan Blue haemocytometer counting after treated with 3 µM MPA or methanol control with the addition of 100 µM guanosine or vehicle control for 3 days. (C) Clonogenicity assay of A375 cells described in A. Clonogenicity was assessed in pre-treated (2 and 3 days) cells seeded at 2000 cells in 6-well plate followed by crystal violet staining (0.5% in methanol) after incubation for 10 days in drug-free media. Representative images after crystal violet-stained wells were shown. (D) 3D matrigel clonogenicity assay of A375 and B16-F10 cells treated with 3 µM MPA or methanol (control) for 10 days. (E) Matrigel-coated Boyden chamber invasion assay of cells described in B. (F) Cell growth analysis done by Trypan Blue haemocytometer counting after A375 cells were treated with siIMPDH2 or scramble control with the addition of 100 µM guanosine or vehicle control for 3 days. (G) Matrigel-coated Boyden chamber invasion assay of cells described in F. (H) Nascent transcripts of the indicated genes were analyzed by qRT-PCR after A375 cells were treated with 3 µM MPA for 0h, 4h and 8h. (I) IF staining with anti-NCL antibody in A375 cells treated with 25 µM Ribavirin, 3 µM MPA, or methanol (Control) with the addition of 100 µM guanosine or vehicle control for 24h. DAPI stains the nuclei. Scale bar: 20 μm (J) RhoA activity assay in A375 cells treated with 3 µM MPA or methanol (Control) with the addition of 100 µM guanosine for 24h. (K) IF staining of A375 cells described in J with anti-p-MLC2 (green) and phalloidin (red). DAPI stains the nuclei. Scale bar: 20 μm. % peripheral p-MLC2 positive cells were plotted on the right plot. Data for A, B, F, H, J and K are from three independent experiments and are presented as mean ±SD, analyzed by one-way or two-way ANOVA plus Tukey’s multiple comparison test. ns: non-significant.

IMPDH2-dependent GTP biosynthesis was shown to support rRNA and tRNA synthesis^7^. Many tumor cells exhibit increased Pol I activity^7, 56–58^, and GTP-dependent Pol I activation has been shown in several cancers^59, 60^. We thus examined RNA synthesis by qRT-PCR in A375 cells and found that both MPA treatment and IMPDH2 silencing blunted pre-rRNA (Pol I transcript), and pre-tRNAl13 (Pol III transcript) expression levels, but not pre-GAPDH mRNA (Pol II transcript), in a time dependent manner (Figure 4G and S4I). This correlated with triggering nucleolar stress responses characterized by delocalization of nucleolin and induction of p53 activity (Figure 4H, 4I, S4J, S4K and S4L), with both effects reversed by guanosine (Figure 4H, 4I). We thus conclude that IMPDH2 regulates ribosome biogenesis in melanoma cells via *de novo* GTP synthesis.

GTP is also essential for G-protein activity^11^, and Rho-GTPases regulate the actomyosin cytoskeleton via ROCK I/II activation and phosphorylation of MLC2 in myosin II to promote melanoma progression^3, 4, 61^. MPA treatment and IMPDH2 silencing in melanoma cells reduced RhoA activity and phospho-MLC2/F-actin levels (Figure 4J and S4M), and guanosine again rescued these effects (Figure 4J and S4M), indicating that IMPDH2 regulates actomyosin contractility via GTP synthesis.

### EZH2 promotes clonogenic and invasive phenotypes via IMPDH2 and cellular GTP

IMPDH2 is the rate-limiting enzyme in the production of GTP^62–64^. Because EZH2 regulated IMPDH2 tetramerization and activity (Figures 3M and 3N), we checked the contribution of EZH2 to cellular GTP production in melanoma. Stable EZH2 knockdown reduced GTP levels by 50% in A375 cells, an effect that was reversed by overexpression with N-terminal domain of EZH2 (1-340) [clone 2] (Figure 5A), which we previously identified as an interaction site for IMPDH2 (Figures 3D-F). Concurrently, EZH2 knockdown also reduced cell proliferation, migration and invasion in a guanosine-reversible manner (Figure 5B and S5A-D). Moreover, EZH2-WT or - H689A overexpression induced cell proliferation and invasion, which were reduced to shEZH2 levels by IMPDH2 silencing; again, these effects were rescued by guanosine. These data implicate IMPDH2 as a key intermediary between EZH2 and GTP synthesis in melanoma (Figure 5C, 5D and S5C-D), in which case EZH2 would be expected to modulate critical, IMPDH2- and GTP-dependent functions in cancer cells, such as rRNA metabolism and GTPase activity (Figures 4G-J).

**Figure 5.**
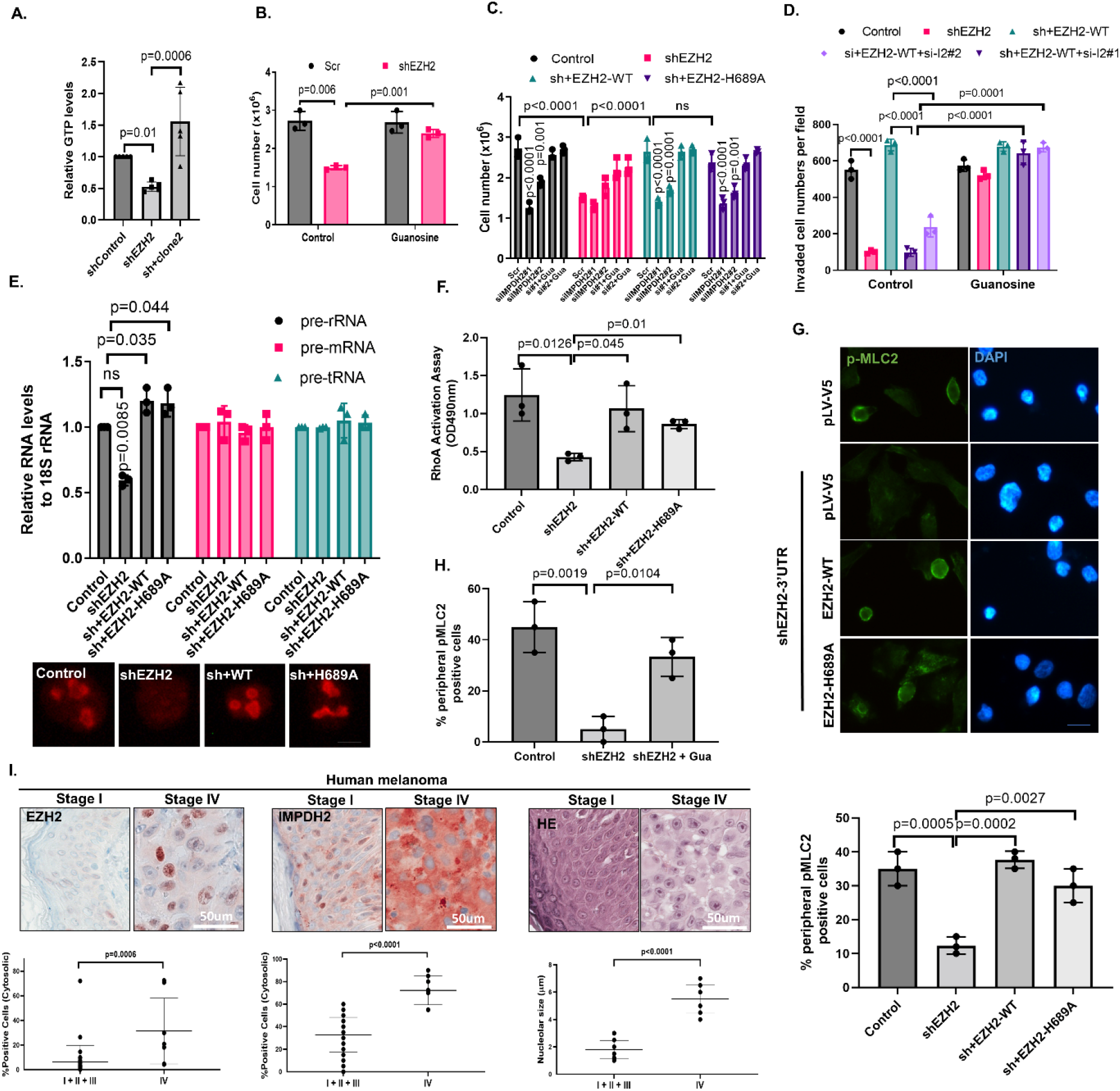
EZH2 regulates clonogenicity/ invasion by regulating rRNA metabolism and Rho GTPase activity via GTP production in melanoma. (A) Relative GTP levels were quantified by HPLC in A375 cells with stable EZH2 knockdown (shEZH2) and overexpression with V5-EZH2-clone2. (B) Cell growth analysis of A375 cells with stable EZH2 knockdown after 3 days of control or 100 µM guanosine addition done by Trypan Blue haemocytometer counting. (C) Cell growth analysis done by Trypan Blue haemocytometer counting and (D) invasion assay counting done by crystal violet staining after A375 cells with stable EZH2 knockdown were rescued by V5-tagged WT-EZH2 or methyltransferase deficient H689A-EZH2 overexpression followed by scramble, si-IMPDH2#1, or si-IMPDH2#2 oligos and 100 µM guanosine addition. (E) Nascent transcripts of the indicated genes were analyzed by qRT-PCR, (F) RhoA activity assay, (G) p-MLC2 IF in A375 cells showing EZH2 knockdown after lentiviral transduction with control shRNA (shControl) or 3′ UTR EZH2-targeting shRNA (shEZH2) and rescue with V5-tagged WT-EZH2 or methyltransferase deficient H689A-EZH2. % Peripheral p-MLC2 positive cells were plotted below images. (H) % peripheral p-MLC2 positive cells were plotted. (I) Human melanoma samples from grade I to IV were stained with anti-EZH2 and anti-IMPDH2 antibodies. Grade I, II, III: n=31 grade IV: n=8. Nucleolar sizes were measured from HE stained samples. Scale bar: 50 μm. Data are presented as mean ±SD, analyzed by student t-test. Data for A-H are from three independent experiments and are presented as mean ±SD, analyzed by one-way or two-way ANOVA plus Tukey’s multiple comparison test. ns: non-significant.

Consistent with this, EZH2 knockdown reduced rRNA synthesis and ribosome biogenesis and induces p53 in melanoma cells (Figures 5E, S5E and S5F) in a guanosine-reversible manner (Figure S5E). Additionally, RhoA activity and phospho-MLC2 levels were lowered by EZH2 silencing (Figure 5F, 5G, S5G, S5H and S5I) and similarly restored by guanosine (Figure 5H and S5G). Reduced phospho-MLC2 was also seen using siEZH2 constructs or EZH2 degradation by DZNep or MS1943, but not by use of EZH2 methyltransferase inhibitors GSK126 and EPZ-6438 (Figure S5J, S5K), consistent with a methyltransferase-independent role for EZH2 in RhoA dependent myosin II activation. In summary, EZH2 regulates levels and critical functions of cellular GTP in melanoma via IMPDH2.

### Nucleolar size and the cytosolic localization of EZH2-IMPDH2 interactions are increased during melanoma progression in patients

If cytosolic EZH2-IMPDH2 interactions drive melanoma cell proliferation and invasion as above, then increased cytosolic expression of these proteins might be expected during melanoma progression in patients. To test this, we interrogated the Protein Atlas database (https://www.proteinatlas.org/ ENSG00000178035-IMPDH2/tissue/skin#img), observing IMPDH2 expression but undetectable EZH2 labelling in the nuclei of normal human melanocytes. Consistent with this, in immunostaining of normal human melanocytes and melanoma samples, EZH2 and IMPDH2 expression were either not detectable or nuclear in normal melanocytes and stage I melanoma samples (Figures 5I and S5M). In stage IV metastatic melanomas, however, cytosolic EZH2 and IMPDH2 expression were significantly increased and asociated with increased cellular nucleolar size, an indicator of ribosome biogenesis (Figure 5I). A functional, methyltransferase-independent link between nucleolar size and EZH2 was verified in tumours from A375 cells in which EZH2 was depleted by shEZH2-3’UTR and then re-expressed with either wild-type or methyltransferase-deficient EZH2 (Figure S5L). These data are consistent with the possibility that cytosolic EZH2-IMPDH2 interactions drive ribosome biogenesis during melanoma progression in patients.

### Sappanone A impedes EZH2-IMPDH2 interactions and melanoma progression

As EZH2 interacts with CBS domain of IMPDH2 (Figure 3C), we sought to test drugs that can inhibit this interaction. A small molecule called Sappanone A (SA) was demonstrated specifically to inhibit IMPDH2 by directly targeting the conserved cysteine residue 140 (Cys140) in the CBS domain of IMPDH2, inducing an allosteric effect on the catalytic pocket that suppressed IMPDH2 activity. We thus examined effects of SA on EZH2-IMPDH2 interactions and melanoma progression. SA inhibited both endogenous EZH2-IMPDH2 interactions, EZH2-IMPDH2-CBS domain interactions and IMPDH2 tetramerization in A375 and B16-F10 cells in a dose-dependent manner (Figure 6A-C and S6A-B). In addition, 10 to 20 µM of SA also promoted nuclear localization of IMPDH2 (Figure 6D and S6C). These data indicated that EZH2-IMPDH2 interactions can be targeted by SA.

**Figure 6.**
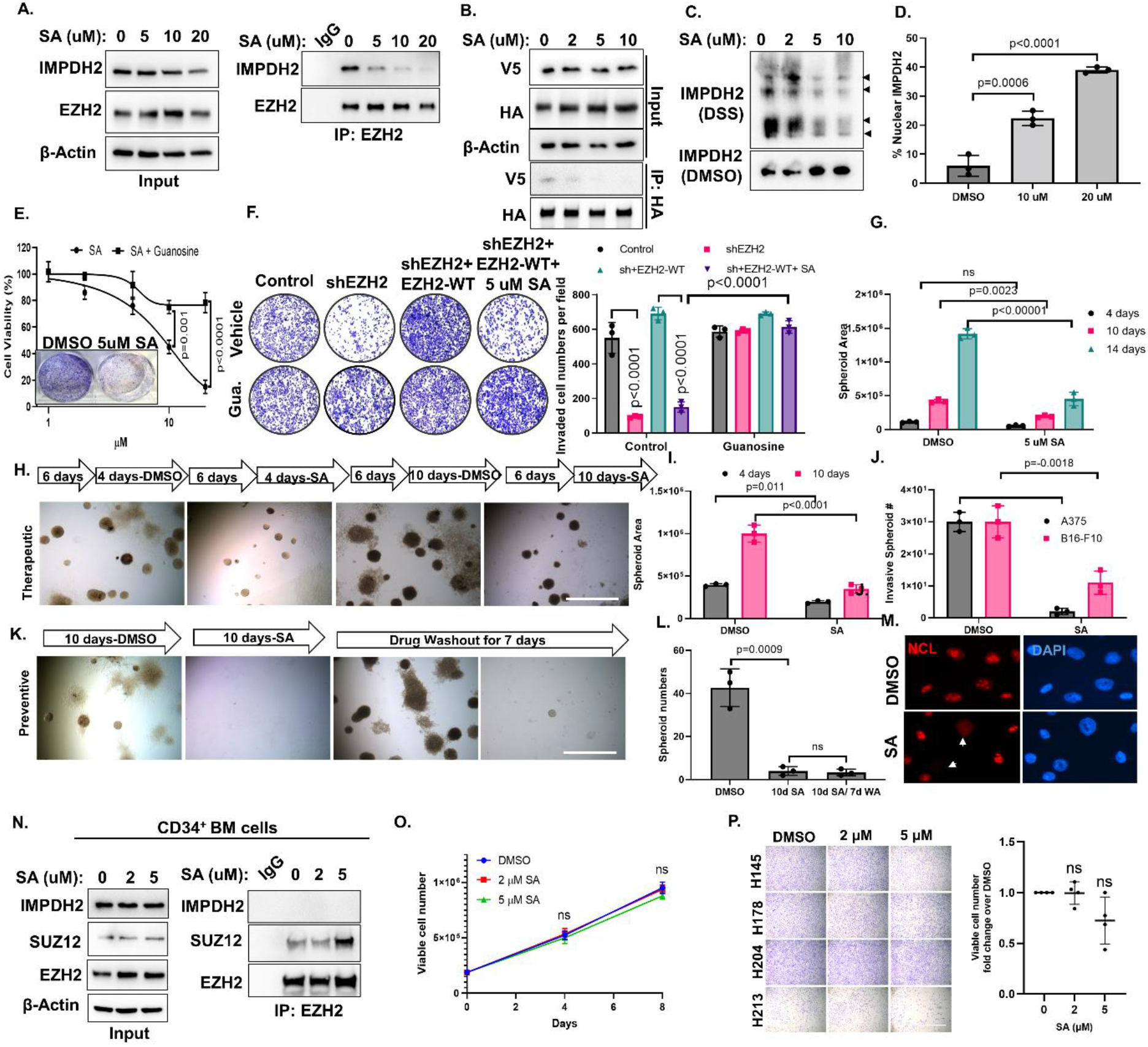
Pharmacological inhibition of EZH2/IMPDH2 interactions by SA attenuates the growth and invasion abilities of melanoma cells *in vitro*. (A) The interaction between endogenous EZH2 and IMPDH2 upon 16h SA treatment (DMSO, 5, 10, 20 µM) was determined in A375 cells by IP with anti-EZH2 antibody followed by WB with anti-EZH2 and anti-IMPDH2 antibody. The inputs were shown on the left. (B) The interaction between HA-EZH2 and V5-IMPDH2-CBS upon 16h SA treatment (DMSO, 2, 5, 10 µM) was determined in A375 cells by IP with anti-HA antibody followed by WB with anti-V5 and anti-HA antibody. The inputs were shown above the IP blots. (C) The clusters of IMPDH2 tetramer were detected from cross-linked whole-cell extracts isolated from A375 cells treated with SA for 16h. (D) Cytosolic versus nuclear localizations of EZH2 and IMPDH2 were examined upon SA treatment in A375 cells by Co-IF. % Nuclear IMPDH2 positive cells were plotted on the group. (E) Dose dependent cell growth curve of A375 cells treated with the indicated dose of SA and - /+ 100 µM guanosine for 3 days. Clonogenicity was shown in the inlet. (F) Matrigel-coated Boyden chamber invasion assay in A375 cells with stable EZH2 knockdown and later rescued by V5-tagged WT-EZH2 overexpression followed by scramble, si-IMPDH2#1, or si-IMPDH2#2 oligos and 100 µM guanosine addition. (left), invaded cell numbers per field were plotted on the right graph. (G) Spheroid areas of 3D colonies grown for 4, 10, 14 days with DMSO or 5 µM SA containing culture medium were measured by Image J program. (H) Sphere formation in 3D Matrigel (therapeutic). A375 cells were grown in Matrigel for 6 days in the absence of SA followed by 4d and 10d days with DMSO or 10 µM SA. Spheroid areas were measured by Image J program and presented in the graph (I). Invasive spheroid numbers were counted manually and presented in the graph (J). (K) Sphere formation in 3D Matrigel (preventive). A375 cells were grown in Matrigel for 10 days in presence of either DMSO (control) or 10 µM SA and then the colonies were grown 7 more days without SA or DMSO and spheres were counted manually and presented in the graph (L). (M) The effect of SA on ribosome biogenesis was measured in A375 cells treated with the indicated doses of SA by anti-NCL antibody. DAPI stains the nuclei. Scale bar: 20 μm. (N) The effect of SA on EZH2 and IMPDH2 interaction was shown by Co-IP coupled WB in CD34^+^ BM cells. Cell growth analysis of CD34+ bone marrow progenitor cells (n=2 patients in triplicates) (O) and normal human melanocytes (n=4) (P) treated with DMSO (vehicle), 2 µM, or 5 µM SA for the indicated time points. Data for D, E, F, G, I, J and L are from three independent experiments and are presented as mean ±SD, analyzed by one-way or two-way ANOVA plus Tukey’s multiple comparison test. ns: non-significant.

*In vitro*, SA reduced melanoma cell proliferation and clonogenicity dose-dependently and in a manner that was reversed by guanosine (Figure 6E and S6D, S6E). Moreover, induction of cell proliferation and invasion by EZH2-WT overexpression following shEZH2 transduction were reduced to those of shEZH2 by 5 µM SA treatment, an effect that was able to be rescued by guanosine (Figure 6F and S6F). In 3D Matrigel spheroid assays, pre-treatment with SA for 10 days prior to seeding spheroid cultures or treatment of established spheroids demonstrated profound anti-melanoma effects, even after 7 days of SA drug washout (Figures 6G-J and S6G-I). SA also induced ribosomal stress and reduced myosin II activation in A375 cells (Figure 6M and S6L).

To examine normal cells that might be susceptible to targeting the EZH2-IMPDH2 interactome, we performed LC/MS on EZH2-immunoprecipitated lysates from CD34^+^ human bone marrow progenitor cells. Although IMPDH2 was detected in anti-EZH2 immunoprecipitates, this could not be verified by CoIP-coupled WB in the same stringency conditions used previously (Table S2 and Figure 6N). Consistent with this, SA has no observable cytotoxic effect on blood progenitors (Figure 6O). Similarly, SA treatment of freshly isolated normal human melanocytes for 7 days at 2 to 5 µM did not significantly attenuate cell growth (Figure 6P). These data indicate that pharmacological inhibition of EZH2-IMPDH2 interactions by SA attenuates melanoma progression without melanocyte and blood cell toxicity by impeding rRNA metabolism and actomyosin contractility.

### EZH2-IMPDH2 interactions and Sappanone A treatment in uveal melanoma and non-melanoma cancers

Although increased EZH2 and IMPDH2 have been linked to many solid cancers^10^, potential interactions between them have not been reported. We thus extended the above studies, examining EZH2 and IMPDH2 levels and interactions in uveal melanoma (92.1 and OMM1), and ovarian (OVCAR-3 and OVCAR-8), breast (MCF7 and MDA-MB-231), and prostate (LNCaP and C4:2) cancer cell lines. Cytosolic EZH2/IMPDH2 interactions were seen in all lines tested (Figures 7A-E and S7A-E). Moreover, total EZH2 degradation by MS1943 treatment, but not treatment with GSK126, a EZH2 methyltransferase inhibitor, attenuated OMM1, OVCAR-8, MDA-MB-231, and C4-2 cell growth. We therefore also tested effects of SA on EZH2-IMPDH2 interactions and IMPDH2 activity in OMM1, OVCAR-8, MDA-MB-231, and PC3 cells, observing reduced EZH2-IMPDH2 interactions (Figure 7F-G, S7F-G) and IMPDH2 tetramerization in every case (Figure S7H-K). These data suggest that cytosolic EZH2-IMPDH2 interactions may be a therapeutic target in a range of cancers beyond cutaneous melanoma.

**Figure 7.**
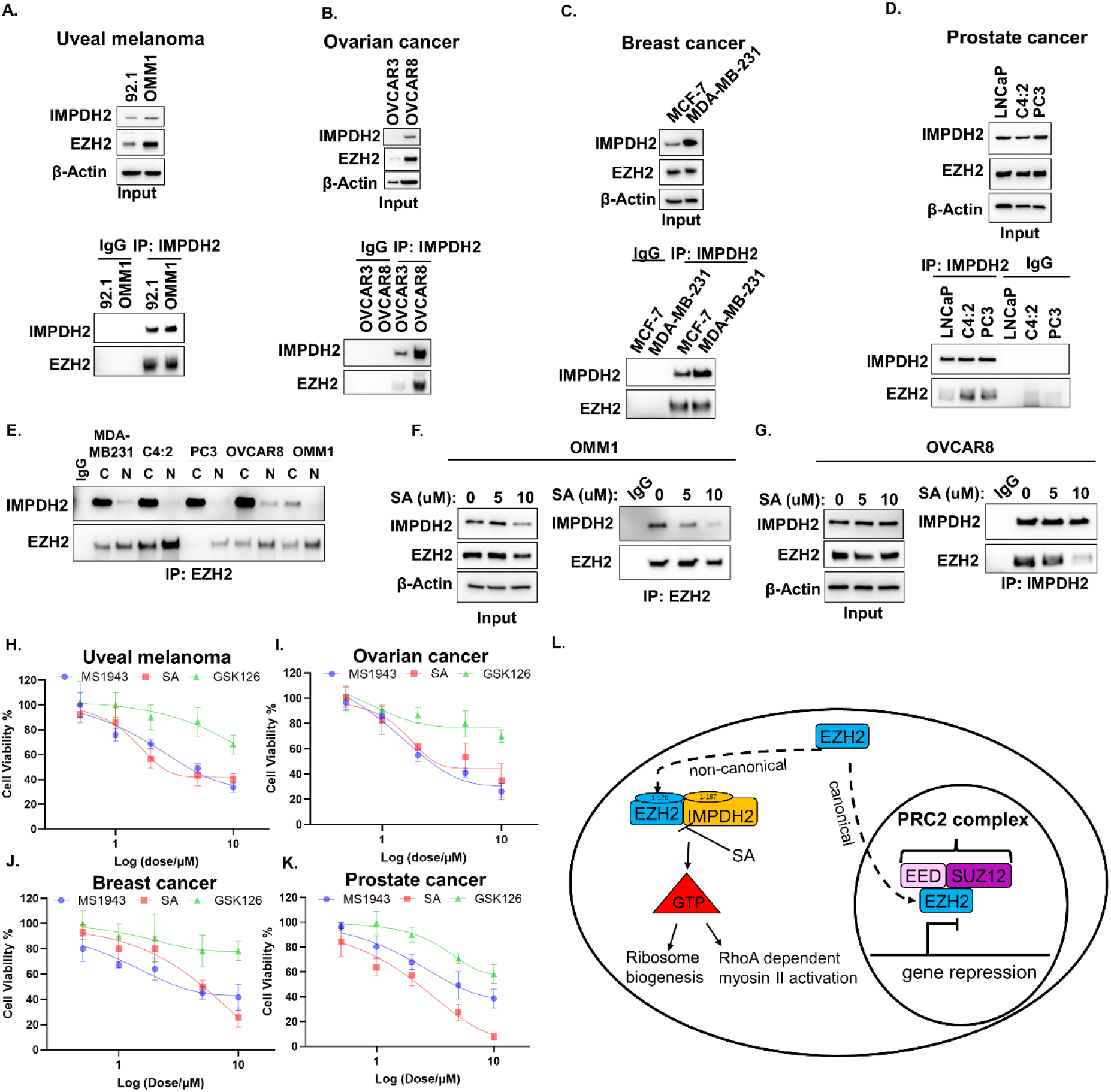
EZH2-IMPDH2 interaction is commonly seen in uveal melanoma, breast, prostate, ovarian cancer, and SA attenuates their growth in *vitro*. EZH2 and IMPDH2 interaction was shown by IP with anti-IMPDH2 antibody followed by WB with anti-EZH2 and anti-IMPDH2 antibody in (A) uveal melanoma, (B) ovarian cancer, (C) breast cancer and (D) prostate cancer cell lines. Inputs were shown at the top of each Co-IP blots. (E) Cytosolic/Nuclear fractionation was done for MDA-MB-231, C4-2, PC3, OVCAR8 and OMM1cells followed by IP with anti-EZH2 antibody followed by western blotting with anti-EZH2 and anti-IMPDH2 antibody. The effect of SA on EZH2 and IMPDH2 interaction was shown by Co-IP coupled WB in (F) OMM1 and (G) OVCAR8 cells. Dose-dependent growth curves of (H) OMM1, (I) OVCAR8, (J) MDA-MB-231 and (K) C4:2 cells upon SA treatment for 3 days. (L) Proposed model depicting both canonical nuclear and non-canonical cytosolic functions of EZH2 as an epigenetic silencer and as GTP regulator via IMPDH2 interaction, which can be blocked by SA. EZH2 induces tumorigenicity and metastasis in melanoma by upregulating rRNA metabolism and RhoA dependent actomyosin contractility via GTP production.

Broadly, our findings support a model of EZH2 function in which its canonical role in the repressive PRC2 complex is complemented in at least some cancers by methyltransferase independent cytosolic EZH2 sequestration and binding to IMPDH2 to activate GTP synthesis and facilitate ribosome biogenesis and actomyosin contractility, thereby promoting cancer progression (Figure 7L).

## DISCUSSION

EZH2 is a bona-fide oncoprotein in cutaneous and uveal melanomas^24–28, 65^, and breast^66^, prostate^67^, and ovarian^68^ cancers, imparting proliferative, migratory, and invasive cancer cell phenotypes. However, the mechanisms through which it imparts these properties are incompletely understood. Here, we reveal a previously unappreciated, methyltransferase-independent function of EZH2 that acts via cytosolic interactions with and activation of IMPDH2 to maintain cellular GTP.

Recently, uveal melanoma cells were shown to be resistant to EZH2 methyltransferase inhibition^69^ unless exposed to supraphysiological doses^70^. Further, triple-negative breast cancer MDA-MB-231cells and castrate-resistant prostate cancer C4-2 and DU145 cells were also reported to be similarly resistant, although sensitive to total EZH2 silencing, suggesting methyltransferase-independent functions of EZH2^38, 67, 71^. Using an unbiased proteomics approach, we uncovered methyltransferase-independent binding partners of EZH2 in cutaneous melanoma to identify IMPDH2 as a critical mediator of the oncogenic effects of EZH2. Furthermore, we showed that EZH2-IMPDH2 interactions are commonly seen and functionally consequential in other cancers, suggesting that their targeting may represent a common molecular target in human cancer.

Previous studies found that EZH2 controls melanoma growth and metastasis through transcriptional repression of distinct tumor suppressors, such as ciliary genes and AMD1 in N-Ras mutant tumors^27, 28^ and also regulates mechanisms of adaptive resistance to immunotherapy^30^. Recently, the combination of EZH2 and MEK inhibition was found to reduce tumor burden markedly in *NRAS* mutant cells, but not *BRAF* mutant cells^72^. This is consistent with our observations of partial anti-melanoma effects following EZH2 methyltransferase inhibition in *NRAS* mutant melanoma cells. In contrast, *BRAF* mutant melanoma cells were resistant to EZH2 methyltransferase inhibition but sensitive to total EZH2 silencing. Methyltransferase-dependent functions of EZH2 may be more prominent in *NRAS* mutant melanomas due to lower expression levels of IMPDH2 and EZH2 in *NRAS* mutant cell lines compared with *BRAF* mutant cells.

Cytosolic localization of EZH2 contributes to pro-metastatic behaviors (i.e. invasion, migration), but this may involve methyltransferase dependent or independent mechanisms in different cell types^34^. Cytoplasmic localization of EZH2 was observed in murine fibroblasts, where it retained methyltransferase activity and regulated actin polymerization^36^. In leukocytes, EZH2 methylated the cytoplasmic protein talin-1 to enhance cell migration by inhibiting binding of talin-1 to F-actin^73^. p38-dependent phosphorylation of EZH2 at the T367 residue was shown to induce cytosolic localization of EZH2 in breast cancer cells, where it interacted with cytoskeletal proteins to promotes metastasis^38^. Cytoplasmic EZH2 expression has also been observed in prostate cancer cells^37^. In this study, we identified EZH2 as a regulator of RhoAGTPase activity and actomyosin contractility via RhoA/ROCK/myosin II activation^3, 4, 61, 74^. Consistent with a role for EZH2 in metastasis, we observed cytosolic localization of EZH2 in melanoma cells particularly in association with more advanced/metastatic disease. However, this was not explained by differential phosphorylation of EZH2. The mechanism of nuclear-cytosolic EZH2 shuttling in melanoma cells remains to be elucidated, although our identification by LC-MS of interactions between EZH2 and kinesin family protein components suggests a role for the latter.

Our data suggest that by altering the subcellular localization of IMPDH2, cytosolic EZH2 may switch differentiation-inducing nuclear IMPDH2 into proliferation-inducing cytosolic IMPDH2. Nuclear IMPDH accumulates during the G2 phase of the cell cycle or following replicative/oxidative stress, and binds to single-stranded, CT-rich DNA sequences via its CBS domain^75–79^. Thus, in nuclei, IMPDH acts as a transcriptional regulator of histones and E2F genes independently of its enzymatic activity^79^. Interestingly, in mouse melanoma cells, Impdh2 disruption by SA reduced pigmentation via repression of tyrosinase gene expression^80^.

In this study, we confirmed that EZH2 alters the subcellular localization of IMPDH, as EZH2 knockdown enhanced nuclear localization of IMPDH2. Functionally, gene expression analysis showed that pigmentation-related genes (*Tyr, Oca2, Trp1*) were upregulated by Ezh2 knockdown independent of its methyltransferase function. These lines of evidence suggest that EZH2 may regulate pigmentation-related gene expression via regulation of IMPDH2 nuclear localization. We cannot exclude the possibility that EZH2 manipulation may induce oxidative stress that induces nuclear IMPDH2 localization in melanoma cells. However, in normal melanocytes where EZH2 was not observed, we observed nuclear IMPDH2. Thus, in the absence of EZH2, nuclear IMPDH2 may stabilize melanocyte differentiation in normal melanocytes and melanomas, whereas during melanoma progression, augmented cytosolic EZH2 may move IMPDH2 to the cytosol where its GTP-producing enzymatic activity supports cell growth and invasion.

Pharmacological targeting of IMPDH2 may represent a tolerable therapeutic strategy in EZH2-IMPDH2 activated cancers. Although trials of pan-IMPDH inhibitors, such as MPA, tiazofurin and benzamide riboside, have been conducted in patients with leukemia and multiple myeloma^81–84^, these studies were terminated due to neurotoxic side effects^52, 85, 86^. Because IMPDH2 is mainly expressed in rapidly proliferating immunocytes, in contrast to the IMPDH1 isoform in normal human leukocytes and lymphocytes^87, 88^, MPA was shown recently to have more hematological side effects than the IMPDH2 specific inhibitor, SA, *in vivo*^16^. Consistent with this, we demonstrated low or no EZH2-IMPDH2 interactions in CD34^+^ human blood progenitors and SA demonstrated no anti-proliferative effects on these cells, in contrast to melanoma cells. Although we found that MPA is anti-tumorigenic and anti-metastatic in melanoma cells, IMPDH2 specific inhibitors are likely to be better and better tolerated treatment options.

In conclusion, we report a role for the EZH2 oncoprotein in promoting tumorigenesis and metastasis in melanoma cells by interacting with IMPDH2 to promote GTP generation, and thereby mechanisms such as upregulating rRNA metabolism and actomyosin contractility that support cancer progression. Discovery of this previously unappreciated, non-enzymatic, GTP-dependent function of EZH2 opens new avenues for EZH2-targeted therapeutics.

---

**Figure S1.**
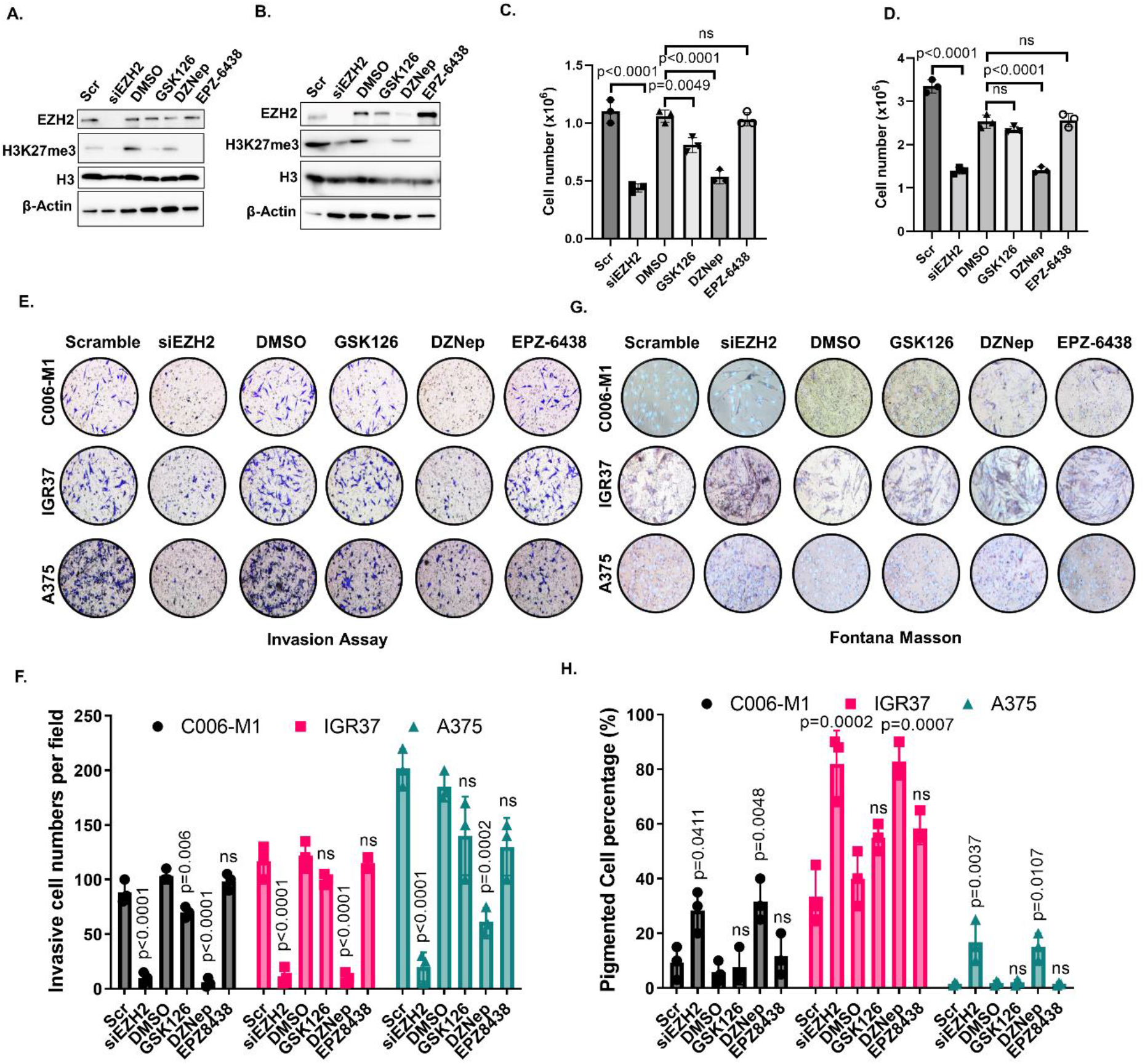
Pharmacological inhibition of EZH2 abundance, but not its activity reduces melanoma cell growth/ invasion and induces pigmentation. C006-M1 and IGR37 cells were treated with siEZH2, 2 μM DZNep, 2 μM GSK126, 2 μM EPZ6438 and scramble or DMSO (control) for 3 days prior to: (A, B) Western blot analysis of EZH2, H3K27me3, H3 and β-Actin protein level, (C, D) cell growth analysis done by Trypan Blue haemocytometer counting, (E) Matrigel-coated Boyden chamber invasion assay was assessed in pre-treated (3 days) cells seeded at 100,000 cells in 24-well plate coated with matrigel after incubation for 24h. (F) Invaded cell counts per well were done by CV staining. (G) Cell pigmentation was assessed by Fontana Masson staining. (H) Pigmented cell percentages were calculated per well. Data for C, D, F and H are from three independent experiments and are presented as mean ±SD, analyzed by one-way ANOVA plus Tukey’s multiple comparison test. ns: non-significant.

**Figure S2.**
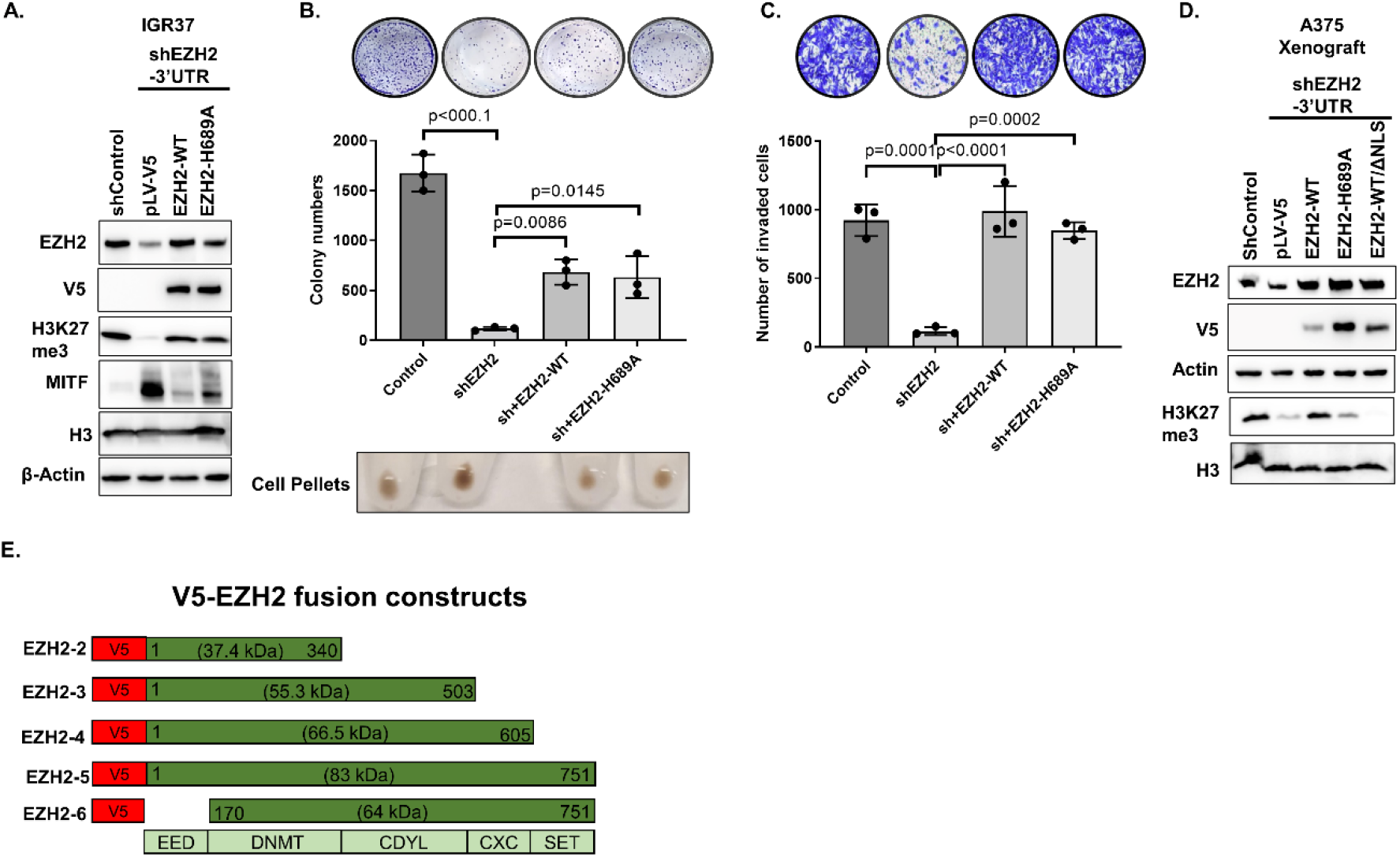
EZH2 has methyltransferase independent function in melanoma clonogenicity, invasion and pigmentation. (A) Western blot analysis of IGR37 cells showing EZH2 knockdown after lentiviral transduction with control shRNA (shControl) or 3′ UTR EZH2-targeting shRNA (shEZH2) and rescue with V5-tagged WT-EZH2 or methyltransferase deficient H689A-EZH2. (B) Clonogenicity assay of cells described in A. Representative images after crystal violet-stained wells were shown above bars and representative images of cell pellets were shown below bars. (C) Matrigel-coated Boyden chamber invasion assay of cells described in A. Representative images after crystal violet staining were shown above bars. (D). Western blot analysis of EZH2, V5, H3K27me3, H3 and β-Actin from A375 xenograft tumor lysates. (E) V5-EZH2 deletion mutant constructs. Data for B, C are from three independent experiments and are presented as mean ±SD, analyzed by one-way ANOVA plus Tukey’s multiple comparison test.

**Figure S3.**
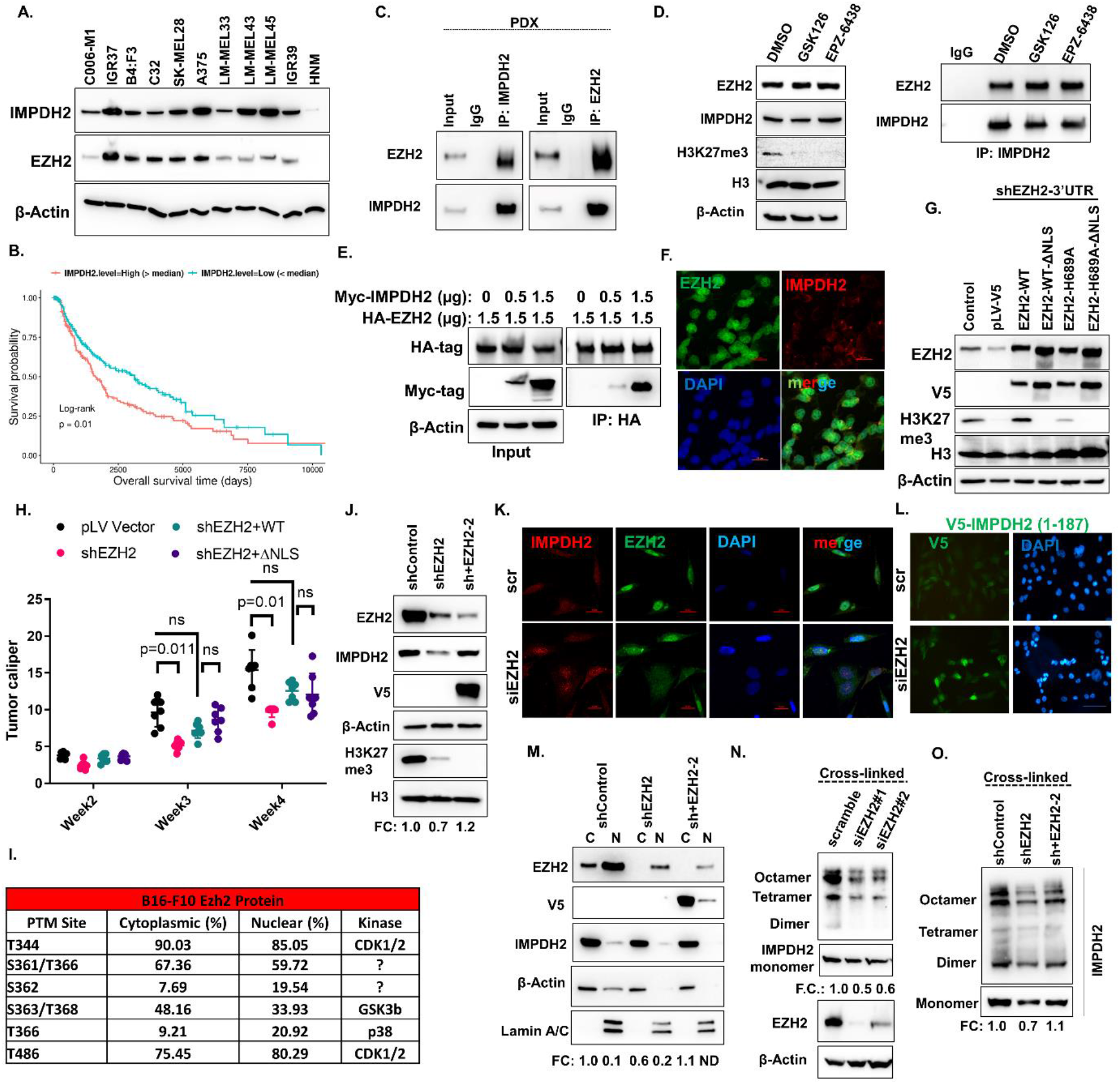
EZH2 interacts with IMPDH2 and induces its tetramerization methyltransferase independently. (A) Western blot analysis of EZH2, IMPDH2 and β-Actin in C006-M1 (NRASQ61K), IGR37 (BRAFV600E), LM-MEL28: B4:F3 (BRAFV600E), C32 (BRAFV600E), SK-MEL28 (BRAFV600E), A375 (BRAFV600E), LM-MEL33 (BRAFV600E), LM-MEL45 (BRAFV600E), IGR39 (BRAFV600E) melanoma cells and normal human melanocytes (NHM). (B) Kaplan-Meier curves of overall survival of TCGA PanCancer Atlas cutaneous melanoma patients (n = 392 patients), stratified by IMPDH2 mRNA levels. Data were analyzed by log rank test. (C) The interaction between endogenous EZH2 and IMPDH2 was determined in PDX tumor lysates by IP with anti-IMPDH2 and anti-EZH2 antibody followed by WB with anti-EZH2 and anti-IMPDH2 antibody. (D) A375 cells were treated with DMSO (control), 2 μM GSK126 or 2 μM EPZ6438 for 2 days prior to: (A) Western blot analysis of EZH2, H3K27me3, H3 and β-Actin (left) and interaction of EZH2/ IMPDH2 were shown by IP with anti-IMPDH2 antibody followed by WB with EZH2 antibody. (E) HA-tagged EZH2-WT and MYC/FLAG-tagged IMPDH2-WT were co-expressed in HEK293 cells. The interaction between overexpressed EZH2 and IMPDH2 was determined by immunoprecipitation with anti-HA antibody followed by western blotting with anti-Myc-tag antibody. (F) Co-immunofluorescence (Co-IF) staining with anti-EZH2 and anti-IMPDH2 antibodies. DAPI stains the nuclei. Scale bar: 20 μm (G) Western blot analysis of EZH2, V5, H3K27me3, H3 and β-Actin in A375 cells described in Figure 3F. (H) Tumor calipers of indicated A375 xenografts (n=7) at the end point. Data are presented as mean ±SD, analyzed by two-way ANOVA plus Tukey’s multiple comparison test. ns: non-significant. (I) Cytosolic and nuclear Ezh2 phosphorylation sites and their percentages measured by LC-MS. Known kinases for the corresponding phospho-sites were also included on the last column of the table.?: Unknown kinases. (J) Western blot of of EZH2, V5, H3K27me3, H3 and β-Actin in A375 cells with control shRNA (shControl) or 3′ UTR EZH2-targeting shRNA and V5-EZH2-clone2. (K) Co-IF staining with anti-EZH2 and anti-IMPDH2 antibodies in C006-M1 cells treated with scramble control or siEZH2 for 3 days. DAPI stains the nuclei. Scale bar: 20 μm (L) IF staining with anti-V5 antibody in A375 cells harboring V5-IMPDH2 (1-187) that was treated with scramble control or siEZH2 for 3 days. DAPI stains the nuclei. Scale bar: 20 μm. (M) Cytosolic/Nuclear fractionation was done from A375 cells shown in J. Lamin A/C is nuclear, and β-Actin is cytosolic marker. The clusters of IMPDH2 tetramer were measured after cross-linking of B16-F10 cells (N) treated with or scramble control, siEzh2#1 orsiEzh2#2 for 3 days and of A375 cells shown in J (O).

**Figure S4.**
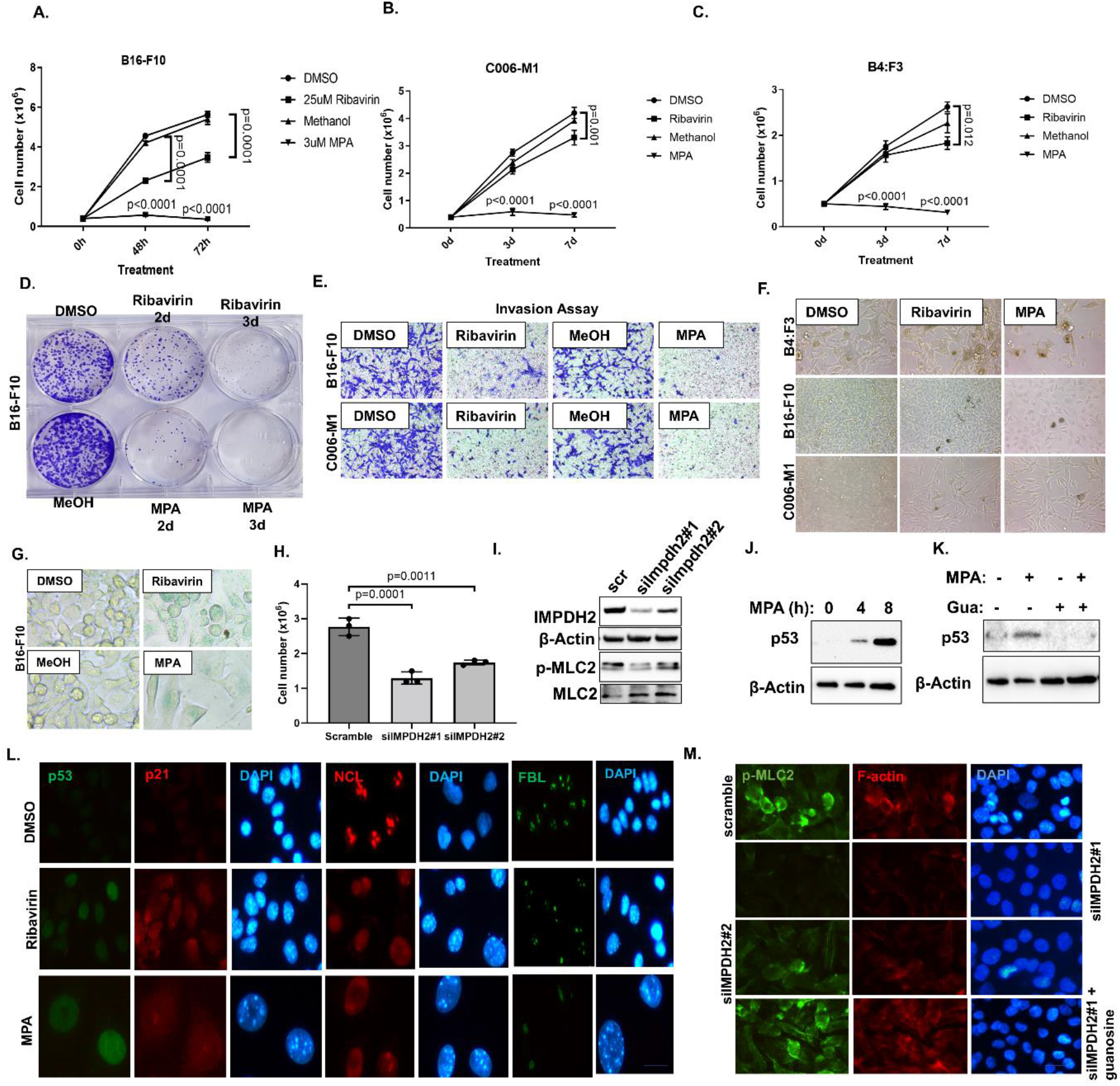
Pharmacological or genetic inhibition of IMPDH2 reduces clonogenicity, invasion by p53 induction and ROCK-myosin II pathway activation. Time-dependent growth curves of A375 (A), C006-M1 (B), B4:F3 (C) cells upon 25 µM Ribavirin or DMSO (control); 3 µM MPA, or methanol (Control). (D) Clonogenicity assay of B16-F10 cells described in A. (E) Matrigel-coated Boyden chamber invasion assay of cells described in A, B. (F) Bright field images of cells described in A, B, C. (G) B16-F10 cell senescence determined by β-gal staining (green). (H) Cell growth analysis done by Trypan Blue haemocytometer counting, (I) Western blot analysis of IMPDH2, β-Actin, p-MLC2 and MLC2 in A375 cells after treated with siIMPDH2#1, siIMPDH2#2 or scramble (control) for 3 days. (J, K) Western blot analysis of p53 and β-Actin in A375 cells with 0h, 4h, 8h 3 µM MPA treatment -/+ 100 µM guanosine. (L) Co-IF staining of A375 cells treated with 25 µM Ribavirin or DMSO (control); 3 µM MPA with anti-p53 (green) and anti-p21 (red) or anti-NCL (red), anti-FBL (green). DAPI stains the nuclei. Scale bar: 100 μm. (M) IF staining of A375 cells described in H with anti-p-MLC2 (green) and phalloidin (red). DAPI stains the nuclei. Scale bar: 20 μm. Data for A, B, C and H are from three independent experiments and are presented as mean ±SD, analyzed by one-way ANOVA plus Tukey’s multiple comparison test.

**Figure S5.**
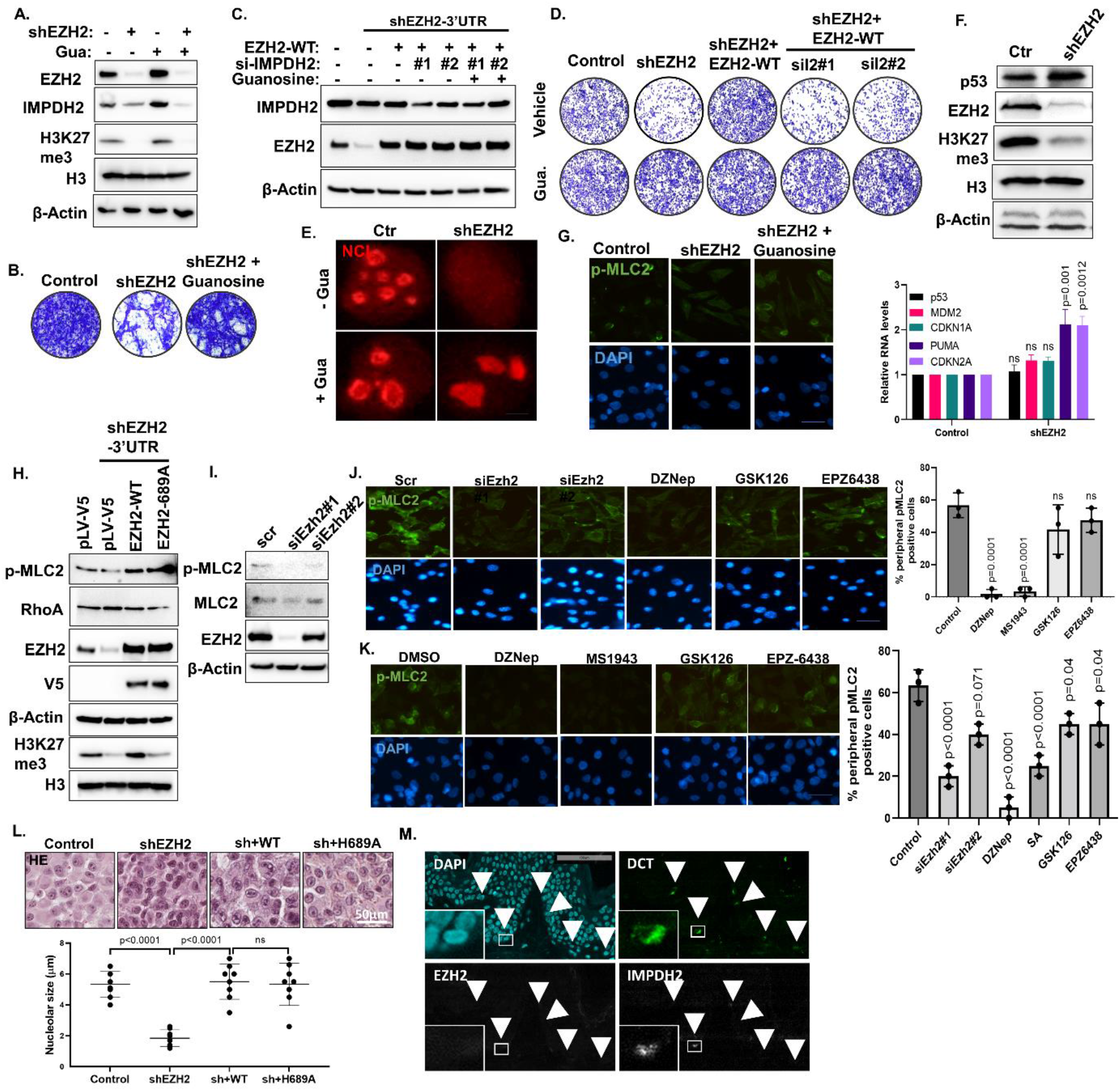
EZH2 modulates ROCK-myosin II activity via Rho GTPase regulation in melanoma cells. (A) Western blot analysis of EZH2, IMPDH2, H3K27me3, H3, β-Actin, (B) Matrigel-coated Boyden chamber invasion assay in A375 cells with stable EZH2 knockdown (shEZH2) -/+ 100 µM guanosine for 3 days. (C) Western blot analysis of IMPDH2, EZH2 and β-Actin and (D) matrigel-coated Boyden chamber invasion assay in A375 cells with stable EZH2 knockdown and rescue with V5-tagged WT-EZH2 with scramble, si-IMPDH2#1, or si-IMPDH2#2 oligos and 100 µM guanosine addition. (E) IF staining with anti-NCL antibody in A375 cells with stable EZH2 knockdown (shEZH2) -/+ 100 µM guanosine for 3 days. (F) Western blot analysis of p53, EZH2, H3K27me3, H3, β-Actin (top) and qRT-PCR of *p53*, *MDM2*, *PUMA*, *CDKN2A* (bottom) in A375 cells with stable EZH2 knockdown. (G) p-MLC2 IF in A375 cells with stable EZH2 knockdown (shEZH2) -/+ 100 µM guanosine for 2 days. (H) Western blot analysis of p-MLC2, RhoA, EZH2, V5, β-Actin, H3K27me3 and H3 in A375 cells showing EZH2 knockdown shRNA (shControl) or 3′ UTR EZH2-targeting shRNA (shEZH2) and rescue with V5-tagged WT-EZH2 or methyltransferase deficient H689A-EZH2. (I) Western blot analysis of p-MLC2, MLC2, EZH2 and β-Actin in B16-F10 cells treated with scramble (control), siEzh2#1, or siEzh2#2 for 3 days. IF staining with anti-p-MLC2 antibody in B16-F10 (J) and A375 cells (K) treated with siEzh2#1, siEzh2#2, 2 µM DZNep, 2 µM GSK126, or 2 µM EPZ6438 for 2 days. DAPI stains the nuclei. Scale bar: 20 μm. % peripheral p-MLC2 positive cells were plotted next to the images. (L) Nucleolar sizes were measured from HE stained xenograft samples (n=7) obtained in Fig. 2F. Scale bar: 50 μm. (M) Co-IF with anti-DCT (melanocyte marker), anti-EZH2 and anti-IMPDH2 in normal human skin samples. Data for F, J, K and L are from three independent experiments and are presented as mean ±SD, analyzed by one-way ANOVA plus Tukey’s multiple comparison test. ns: non-significant.

**Figure S6.**
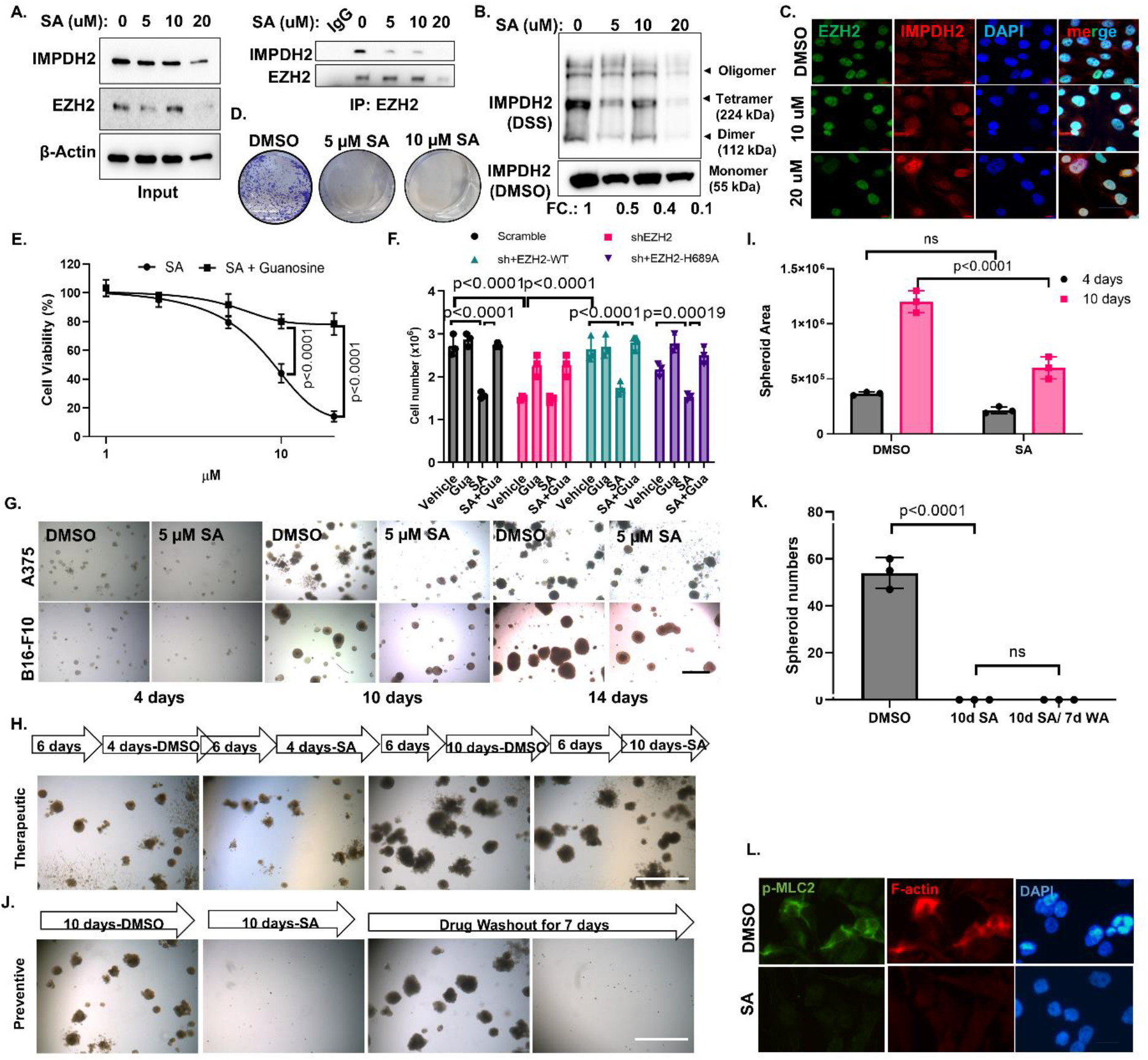
SA reduces EZH2/IMPDH2 interaction, IMPDH2 tetramerization/ nuclear translocation and attenuates the growth and invasion abilities of melanoma cells *in vitro*. (A) The interaction between endogenous EZH2 and IMPDH2 upon 16h SA treatment (DMSO, 5, 10, 20 µM) was determined in B16-F10 cells by Co-IP with anti-EZH2 antibody followed by WB with anti-EZH2 and anti-IMPDH2 antibody. The inputs were shown on the left. (B) The clusters of IMPDH2 tetramer were detected from cross-linked whole-cell extracts isolated from B16-F10 cells treated with indicated dose of SA for 16h. (C) Cytosolic versus nuclear localizations of EZH2 and IMPDH2 were examined upon SA treatment in A375 cells by Co-IF using anti-EZH2 (green) and anti-IMPDH2 (red) antibodies. DAPI stains the nuclei. Scale bar: 20 μm. (D) Clonogenicity assay of A375 cells described in A. (E) Dose dependent cell growth curve of B16-F10 cells treated with the indicated doses of SA and -/+ 100 µM guanosine for 3 days. (F) Cell growth analysis of A375 cells with stable EZH2 knockdown and rescue with V5-tagged WT-EZH2 or methyltransferase deficient H689A-EZH2 followed by DMSO or 5 µM SA and 100 µM guanosine addition. (G) Time dependent sphere formation in 3D Matrigel (preventive). A375 and B16-F10 cells were grown in Matrigel for 4 d, 10 d and 14 d in the presence of either DMSO (control) or 5 µM SA. (H) Sphere formation in 3D Matrigel (therapeutic). B16-F10 cells were grown in Matrigel for 6 days in the absence of SA followed by 4d and 10d days with DMSO or 10 µM SA. Spheroid areas were measured by Image J program and presented in the graph (I). (J) Sphere formation in 3D Matrigel (preventive). B16-F10 cells were grown in Matrigel for 10 days in presence of either DMSO (control) or 10 µM SA and then the colonies were grown 7 more days without SA or DMSO and spheres were counted manually and presented in the graph (K). (L) The effect of SA on actomyosin contractility was measured in A375 cells treated with DMSO or 5 µM SA for 3 days with anti-p-MLC2 antibody (green)/ Phalloidin (red) IF. DAPI stains the nuclei. Scale bar: 20 μm. Data for E, F, I and K are from three independent experiments and are presented as mean ±SD, analyzed by one-way or two-way ANOVA plus Tukey’s multiple comparison test. ns: non-significant.

**Figure S7.**
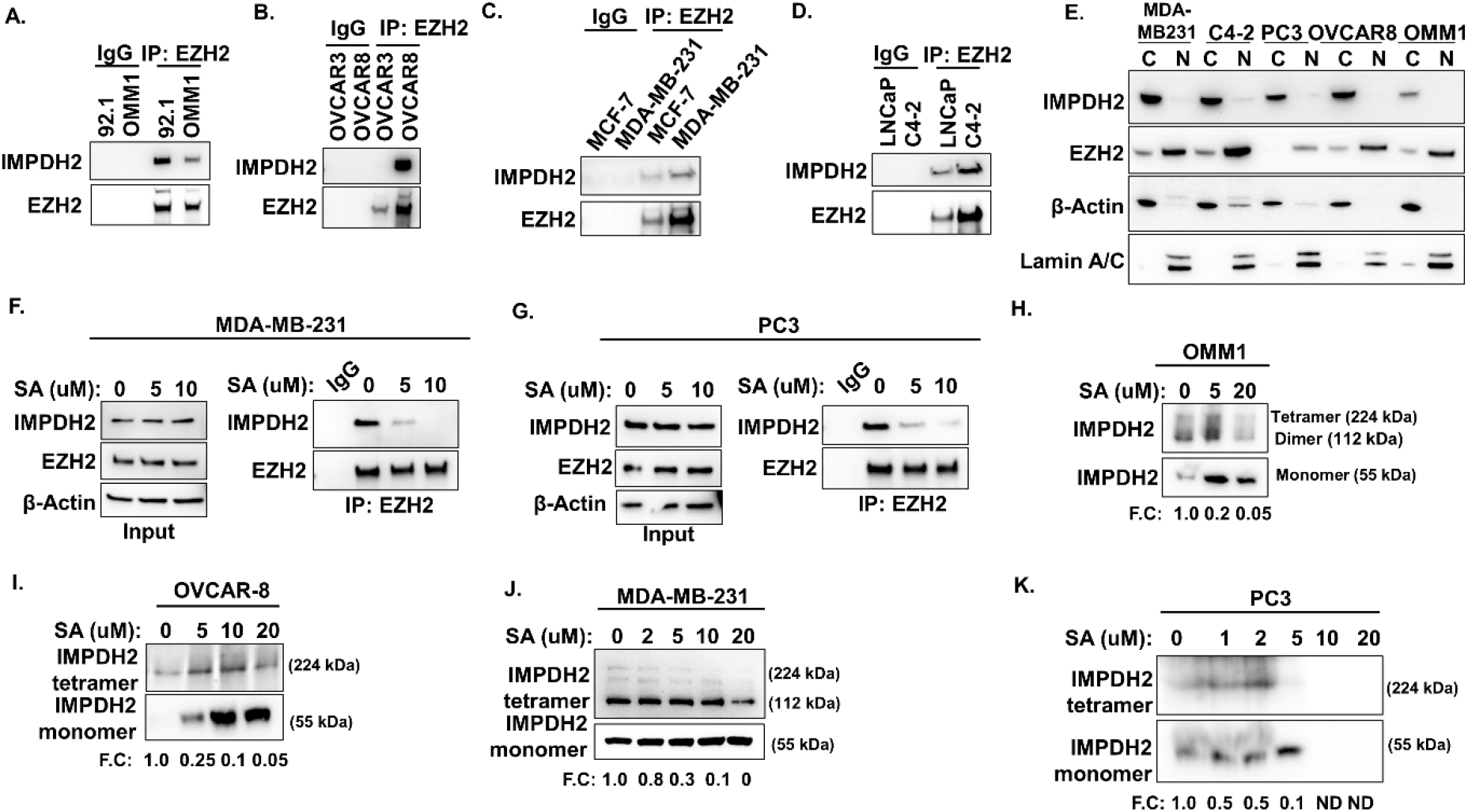
Cytosolic EZH2-IMPDH2 interaction is seen in uveal melanoma, breast, prostate, ovarian cancer, and SA attenuates IMPDH2 tetramerization. EZH2 and IMPDH2 interaction was shown by IP with anti-EZH2 antibody followed by WB with anti-EZH2 and anti-IMPDH2 antibody in (A) uveal melanoma, (B) ovarian cancer, (C) breast cancer and (D) prostate cancer cell lines. (E) Cytosolic/Nuclear fractionation was done for OMM1, OVCAR8, MDA-MD231 and C4:2 cells followed by IP with anti-IMPDH2 antibody followed by western blotting with anti-EZH2 and anti-IMPDH2 antibody. Lamin A/C is nuclear, and β-Actin is cytosolic marker. Inputs were shown above the IP blot. The effect of SA on EZH2 and IMPDH2 interaction was shown by Co-IP coupled WB in (F) MDA-MB-231 and (G) PC3 cells. The clusters of IMPDH2 tetramer were detected from cross-linked whole-cell extracts isolated from (H) OMM1, (I) OVCAR8, (J) MDA-MB-231 and (K) PC3 cells treated with the indicated dose of SA for 16h.

